# Regulating bacterial behavior within hydrogels of tunable viscoelasticity

**DOI:** 10.1101/2022.01.06.475183

**Authors:** Shardul Bhusari, Shrikrishnan Sankaran, Aránzazu del Campo

## Abstract

Engineered living materials (ELMs) are a new class of materials in which living organism incorporated into diffusive matrices uptake a fundamental role in material’s composition and function. Understanding how the spatial confinement in 3D affects the behavior of the embedded cells is crucial to design and predict ELM’s function, regulate and minimize their environmental impact and facilitate their translation into applied materials. This study investigates the growth and metabolic activity of bacteria within an associative hydrogel network (Pluronic-based) with mechanical properties that can be tuned by introducing a variable degree of acrylate crosslinks. Individual bacteria distributed in the hydrogel matrix at low density form functional colonies whose size is controlled by the extent of permanent crosslinks. With increasing stiffness and decreasing plasticity of the matrix, a decrease in colony volumes and an increase in their sphericity is observed. Protein production surprisingly follows a different pattern with higher production yields occurring in networks with intermediate permanent crosslinking degrees. These results demonstrate that, bacterial mechanosensitivity can be used to control and regulate the composition and function of ELMs by thoughtful design of the encapsulating matrix, and by following design criteria with interesting similarities to those developed for 3D culture of mammalian cells.

## 1. Introduction

The combination of synthetic biology and materials science has given rise to the field of engineered living materials (ELMs), wherein live organisms (bacteria, yeast, algae etc.) become active components of material’s design and perform advanced functions.^[1-3]^ Examples of ELMs include biofilters to sequester metals^[4]^ or viruses,^[5]^ bacterial hydrogels for biosensing,^[6]^ shape-morphing composites,^[7]^ self-healing adhesives,^[8]^ photosynthetic bio-garments^[9]^ or self-regulated drug delivery devices.^[10]^ A common feature in these constructs is the encapsulation of the organisms within matrices including natural polymers like agarose,^[11,12]^ alginate,^[13]^ and dextran,^[14]^ synthetic polymers like polyvinyl alcohol^[15]^ and Pluronic,^[16,17]^ or inorganic matrices like porous silica.^[18]^ Alternatively, proteinaceous^[19]^ or cellulose^[6]^ matrices produced by the organisms themselves, as in a biofilm, can serve as encapsulating networks. The matrix confers a protective environment for the cells while it allows diffusion of nutrients and gases to maintain the viability and functionality of the entrapped organisms. It also confines and retains the organisms inside the material, which is a necessary requirement in future application of ELMs containing genetically modified organisms.

Recently, a few studies have highlighted that spatial confinement and matrix mechanical properties affect the growth and functionality of embedded bacteria.^[10,20,21]^ The current understanding, mainly from studies of bacteria or yeast embedded in hydrogels, indicate that the microbes grow inside the hydrogel network to form dense clusters at slower rates than in suspension.^[20,22,23]^ Cell’s response to the mechanical properties of their microenvironment is well known from 3D cultures of mammalian cells, whose proliferation, migration or differentiation programs depend on the viscoelasticity^[24]^ and the degradation kinetics of the hydrogel network.^[25,26]^ Studies in engineered hydrogels with viscoelastic properties that can be modulated by the type of network crosslinks (reversible/dynamic vs. permanent) and by the nature of the degradable sequences have helped to understand and quantify eukaryotic organism’s mechanosensitivity range and response.^[27]^ Preliminary studies from us^[10]^ and others^[21]^ on hydrogel embedded bacteria have indicated that increasing stiffness of the hydrogel network hinders extension of the bacterial cell wall and thus reduces bacterial growth. Based on these observations, we hypothesized that the behavior of encapsulated bacterial colonies, e.g. growth rate or metabolic activity, might be regulated by tuning the viscoelastic properties of the embedding matrix. Hydrogels formed by the triblock copolymer Pluronic F127 (Plu, PEG_106_-PPO_70_-PEG_106_) and its diacrylated derivative (PluDA), previously used for bacterial encapsulation in ELMs, seemed appropriate model systems to test this hypothesis.^[28,29]^ Plu solutions above critical micellar concentration (0.725 wt % at 25 °C ^[30]^) self-assemble in water and form micelles.^[31]^ At concentrations >5 wt% and temperatures above 14°C, micelles self-assemble through physical interactions and form an associative hydrogel (**Figure 1a**). ^[28,29,31]^ Such hydrogels swell and dissociate into individual micelles when immersed in water. When Plu is mixed with PluDA, similar hydrogels are formed but in this case they can be stabilized by covalent cross-linking of the acrylate end-groups. By varying the polymer concentration between 5 and 30 wt%, hydrogels with a storage modulus between 1 and 50 kPa can be obtained.^[31]^

**Figure 1.**
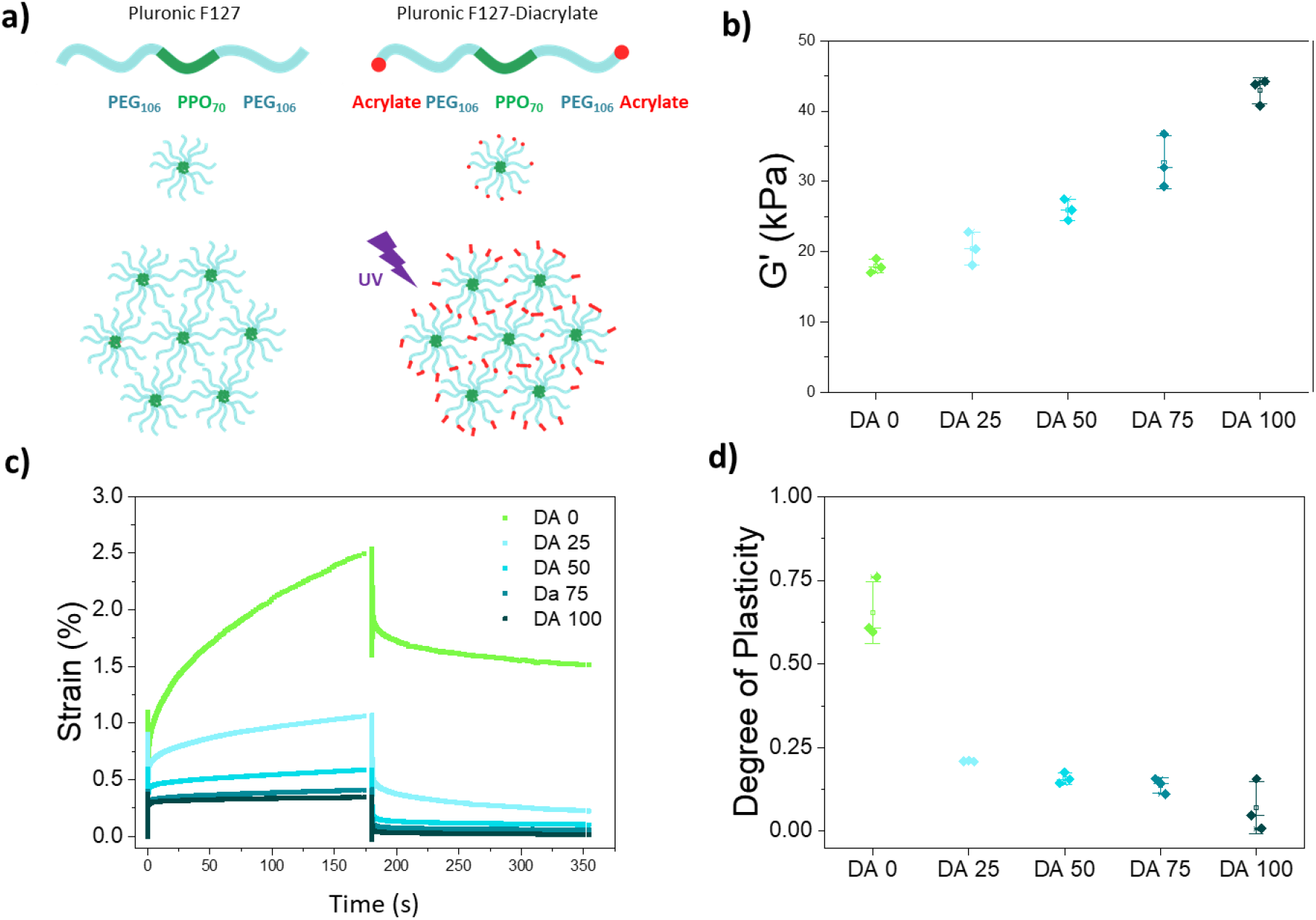
Structure and properties of Plu/PluDA hydrogels (DA 0-100) **a)** The amphiphilic Pluronic F127 chains form micellar assemblies above 14 °C. Acrylate terminated Pluronic-DA introduces covalent inter and intramicellar crosslinks after the photoinitiated radical polymerization step in the presence of Irgacure 2959; **b)** Storage modulus of Plu/PluDA hydrogels with increasing PluDA concentration after photoinitiated polymerization; **c)** Representative creep recovery curves of Plu/PluDA hydrogels and **d)** plasticity values (Residual strain/Peak strain) calculated from **c)**. (N = 3, whiskers indicate standard deviation values and horizontal line in the middle denote median.)

In this work, for the first time, we show that by varying the covalent cross-linking degree in Plu/PluDa hydrogels, we can tune their viscoelastic properties, which in-turn influences the behavior of encapsulated *E. coli*. We reveal fundamental insight on bacterial growth rate and metabolic function in response to material’s mechanics. Our results provide guidance for the design of functional and safe ELMs, by which the function of the embedded organisms is supported, controlled and improved.

## 2. Results

### 2.1. Physicochemical properties of Pluronic F127/Pluronic F127-Diacrylate mixtures

Plu/PluDA hydrogels were prepared by mixing 30% w/v solutions of Plu and PluDA in different ratios (0/100, 25/75, 50/50, 75/25, 100/0) at 4 °C. The 30% w/v Plu/PluDA mixtures formed hydrogels at temperatures above 14°C (**Figure S1g**). To covalently crosslink the hydrogels, a light exposure step was used to photoinitiate the radical polymerization of the acrylate groups. This process yielded transparent hydrogels with constant polymer content, similar organization of the physical network (as reflected by the values of the shear modulus before photopolymerization, Figure S1g), and different degrees of covalent crosslinking.

The combination of physical (reversible) and chemical (permanent) crosslinks confers Plu/PluDA hydrogels (named DA 0 to DA 100 in the following text) viscoelastic properties that vary with PluDA concentration. The hydrogels showed increasing storage modulus (from 18 to 43 kPa) with increasing concentration of PluDA from 0 to 100% (Figure 1b, **Table 1, Figure S2**). In creep-recovery experiments (Figure 1c), DA 0 behaved as a viscoelastic fluid and showed the highest deformation during the loading cycle and the highest residual deformation after recovery. With increasing PluDA ratio, hydrogels became viscoelastic solids and showed smaller deformations and higher elastic recovery. The fluid character of the DA 0 hydrogel is associated with the dynamic nature of the physical network. The polymer chains and the micelles are associated by reversible interactions and can flow easily under the applied load.^[32]^ The covalent bonds introduced by PluDA fix the position of PluDA chains in a permanent crosslinked network and confer elastic properties to the hydrogel. The ratio between the maximum and residual deformations in the creep test is the degree of plasticity (Figure 1d). This value was above 0.6 for DA 0 and decreased to 0.2 in DA 100 hydrogels. In summary, the viscoelastic properties of the Plu/PluDA hydrogels can be tuned depending on the extent of covalent crosslinks incorporated.

**Table 1.**
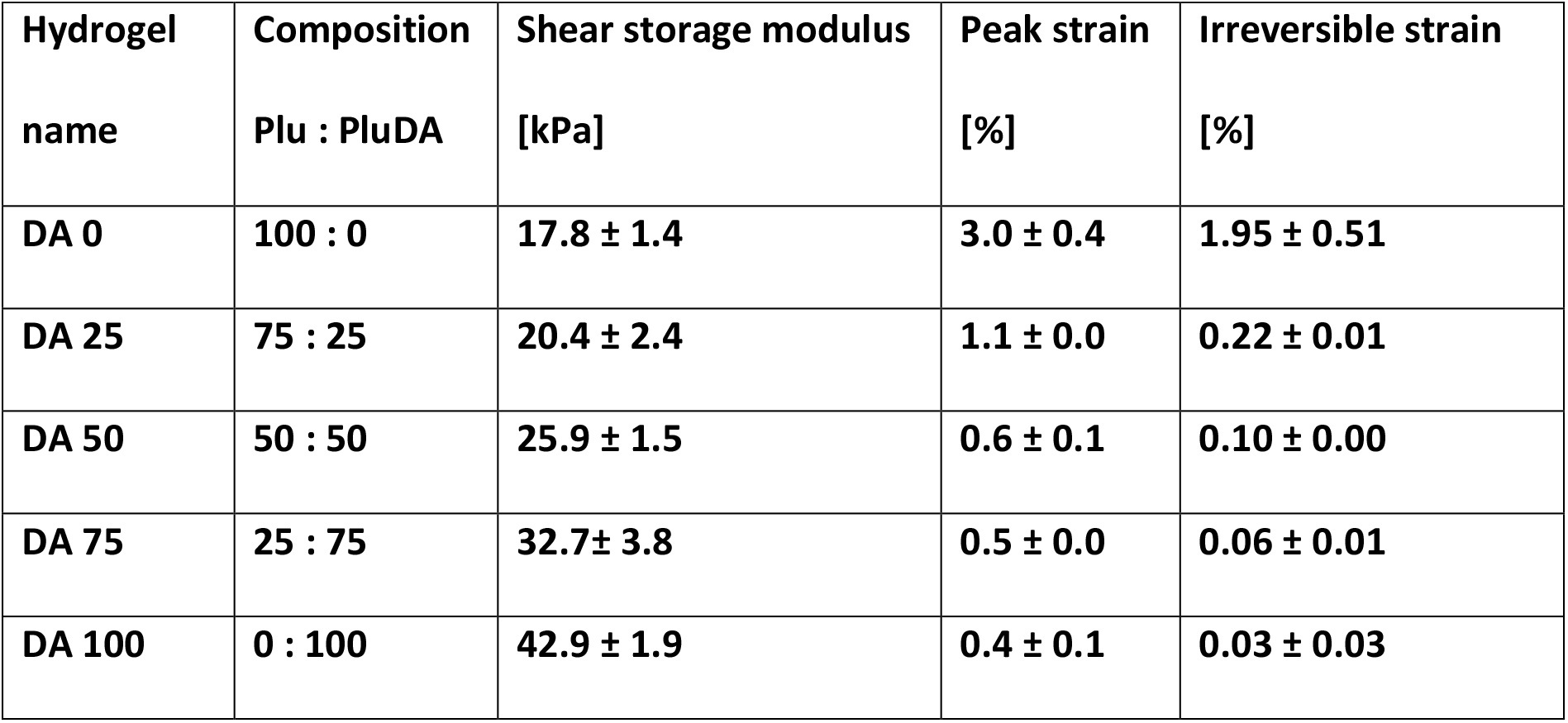
Mechanical properties of DA 0-100 hydrogels with different compositions obtained from rheological analysis: shear storage modulus (G’) of the crosslinked hydrogels (Figure 1b, t = 420 s), maximum deformation (peak strain) at the end of the creep test (Figure 1c, t = 180 s), and the residual deformation (irreversible strain) after recovery (Figure 1c, t = 360 s).

### 2.2. Colony growth and metabolic activity inside Pluronic hydrogels with different ratios of chemical crosslinking

Plu/PluDA hydrogels loaded with bacteria in LB medium were injected in microchannels of a commercial microfluidics chip or formed as films in a microwell plate (**Figure S3**). In both formats bacteria were cultured under static conditions, i.e. no medium exchange. The bacteria laden hydrogels were transparent to the eye and for microscopy. Bacteria encapsulated within microchannels were imaged by bright field microscopy. Right after encapsulation, isolated bacteria were homogenously distributed across the hydrogels (**Figure S4**). Bacteria were not motile inside the hydrogels, presumably due to the physical confinement imposed by the hydrogel network. Bacteria divided inside the hydrogels and daughter cells remained bundled together in what we will hereafter refer to as a ‘colony’. Initially, progeny cells arranged end-to-end, in a chain-like morphology (**Figure 2a-c**). As the length of the colony increased, buckling events occurred along the chain of growing cells, wherein 2 adjacent cells deviated from the linear chain and daughter cells emerging from them formed parallel or branched chains on continued growth (Figure 2a, d). This was observed as an increase in the in-plane width (**Figure S5b**) of the colonies (i.e. measured in the focal plane of imaging), or an increase in contrast when the buckled cells overlapped in the Z direction. Beyond 6 hours, the colony size increased minimally. The rate and extent of colony elongation varied with the ratio of chemical crosslinking in the hydrogel. More pronounced and faster elongation was observed in hydrogels with lower chemical crosslinking ratio (Figure 2b). The overall in-plane length reached by the colonies and the time before their first buckling event occurred decreased with increasing ratio of chemical crosslinking (Figure 2a-d). The mean colony length at 6 h dropped from 13.1 ± 7.1 µm for DA 0 to 1.2 ± 0.6 µm for DA 100 (Figure 2c).

**Figure 2.**
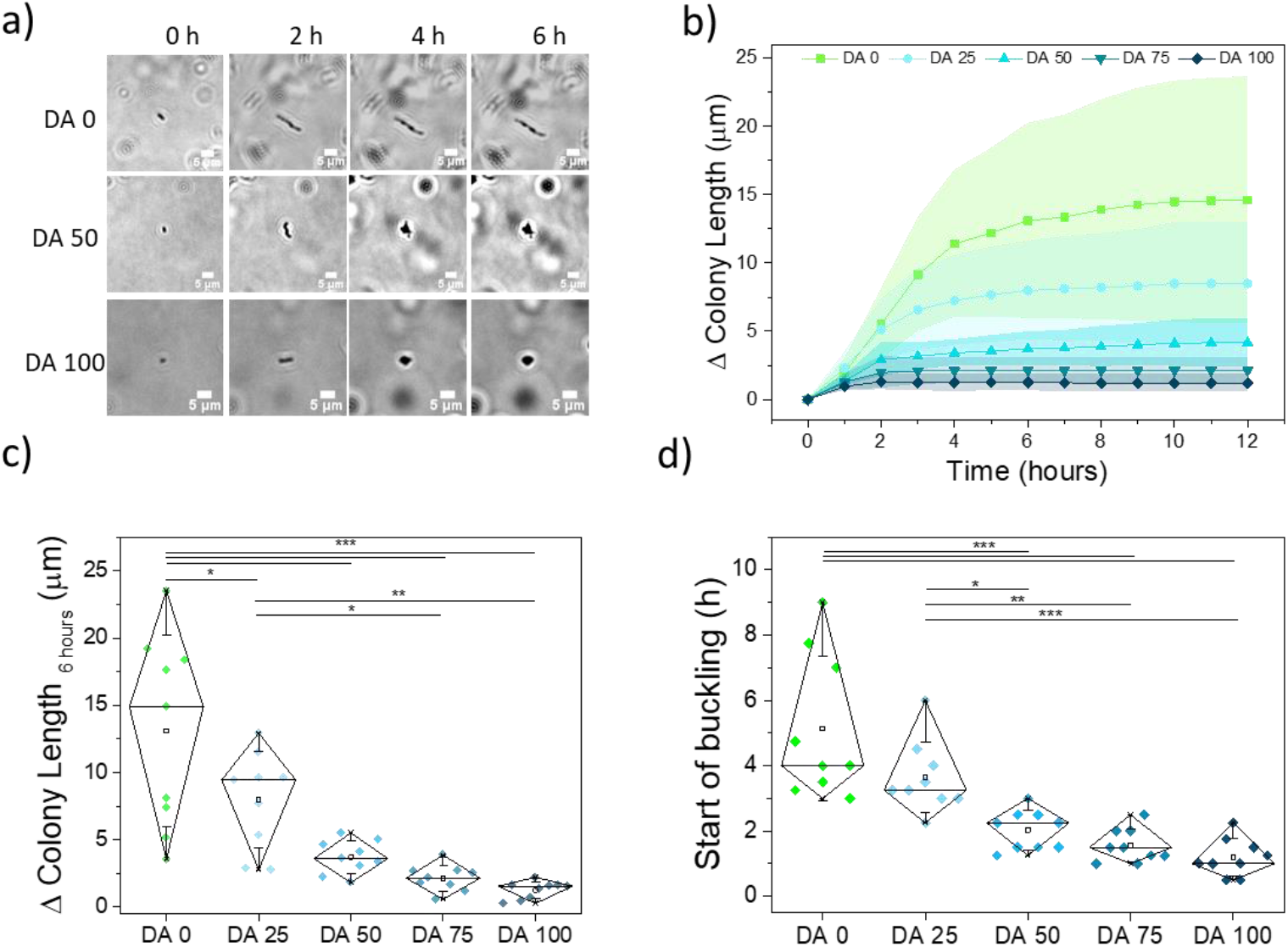
Bacterial growth inside DA 0-100 hydrogels inserted in microchannels, and quantification of geometrical and dimensional features of the growing colonies. **a)** Time lapse bright-field images of bacterial colonies within DA 0, DA 50 and DA 100 hydrogels (scale = 5 μm) representing the differences in the morphology of the growing colonies. **b)** Increase in the in-plane length of the encapsulated bacterial colonies along the longitudinal axis of the first cell with time (mean ± standard deviation). **c)** Increase in the length of individual colonies at 6 h timepoint; **d)** Time point for the first buckling event along the longitudinal axis of a growing bacterial colony in the different hydrogels. N = 9 colonies from 3 independent experiments. (Diamond plots indicate 10 and 90 percentile values, whiskers indicate standard deviation values, * p < 0.05, ** p < 0.01, *** p < 0.001)

The first buckling event occurred at a mean time point of 5.1 ± 2.2 h for DA 0 and at 1.2 ± 0.5 h for DA 100 (Figure 2d). This time point corresponded to the 4^th^ or 10^th^ division cycles in DA 25 or DA0 gels and to the 1^st^ or 2^nd^ division cycle in DA 100. The growing colonies in hydrogels with higher PluDA ratio showed a more rounded shape. This was reflected in the smaller in-plane aspect ratio of the colony (Figure 2a, S5c, f). DA 25, DA 50 and DA 75 showed intermediate behavior with regards to all the above-mentioned aspects. These results indicate that the covalent crosslinking of the hydrogel matrix impacts the growth behavior of the encapsulated bacteria in the 3D network. The higher the chemical crosslinking degree of the matrix, the slower the bacteria elongation rate, the smaller the size of the colonies (as measured within the focal plane), and the more rounded shape the colonies adopt.

Interestingly, the distribution of the in-plane colony length values and of the buckling times narrowed as the ratio of chemical crosslinks in the hydrogel increased, i.e. the permanent crosslinks homogenized the morphology of the individual colonies.

A number of control experiments were performed to assess the impact of nutrients in the observed behavior of encapsulated bacteria. No differences in the diffusion rate of small molecules across the hydrogels with different crosslinking degree was observed in FRAP experiments **(Figure S6)**. Growth experiments performed in hydrogels with a two-fold nutrient concentration showed similar shape of the growth curves and a higher in-plane growth length in DA0 and DA25 hydrogels (**Figure S7**). These results suggest that confinement is the dominant factor regulating growth behavior of *E. coli* in hydrogels with higher chemical crosslinking degree (DA 50-100), whereas bacterial growth in hydrogels with lower chemical crosslinking degrees is also influenced by the nutrient concentration under our experimental conditions. Overall, our data confirm our hypothesis that bacterial growth in hydrogels can be regulated by tuning the design of the encapsulating network, more specifically through the type and degree of crosslinking.

To better investigate the morphology of the colonies in 3D and quantify colony volumes in the different conditions, we used confocal fluorescence microscopy to follow the colony growth. Experiments were performed in hydrogel films within a microwell plate sealed with silicone oil to prevent evaporation of water from the hydrogel while allowing O_2_ diffusion. Since nuclear staining can cause genetic modifications in the bacteria and impact growth behavior, we opted to use a genetically engineered strain of *E. coli* that constitutively produces the fluorescent iLOV protein.^[33]^ This strain grows only slightly slower than unmodified *E. coli* **(Figure S8)**, presumably as consequence of the metabolic burden of expressing the protein. Z-stack images of the bacteria in the hydrogels were acquired at 0, 3, 6 and 24 h as shown in **Figure 3** and **Figure S9**. Under these conditions, isolated bacteria at 0 h grew into colonies of similar morphologies as observed with unmodified *E. coli* in the microchannels (Figure 2a, Figure S4). Bacteria in hydrogels with higher chemical crosslinking degrees formed smaller colonies, in agreement with our observations in the in-plane colony length extension analysis (Figure 2), with mean colony volumes ranging from 472 ± 439 µm^3^ for DA 0 to 213 ± 94 µm^3^ for DA 100 (Figure 3b). Higher chemical crosslinking degrees restricted the maximum size the colonies could reach. The volumes of the larger colonies (95^th^ percentile) drop from 1115 µm^3^ in DA 0 to 359 µm^3^ in DA 100 (Figure 3). Colony volume values were more homogenous (narrower distribution) in gels with higher degrees of chemical crosslinking (Figure 3b). At whole culture level, the volume fraction of the bacteria within the gels increased with time and decreased with DA content (from 0.06% at the start to 2% for DA 0 and to 1.1% for DA 100 by 24 h) (Figure 3c, Figure S9). The morphology of the colonies imaged in 3D was also different across the hydrogels.

**Figure 3.**
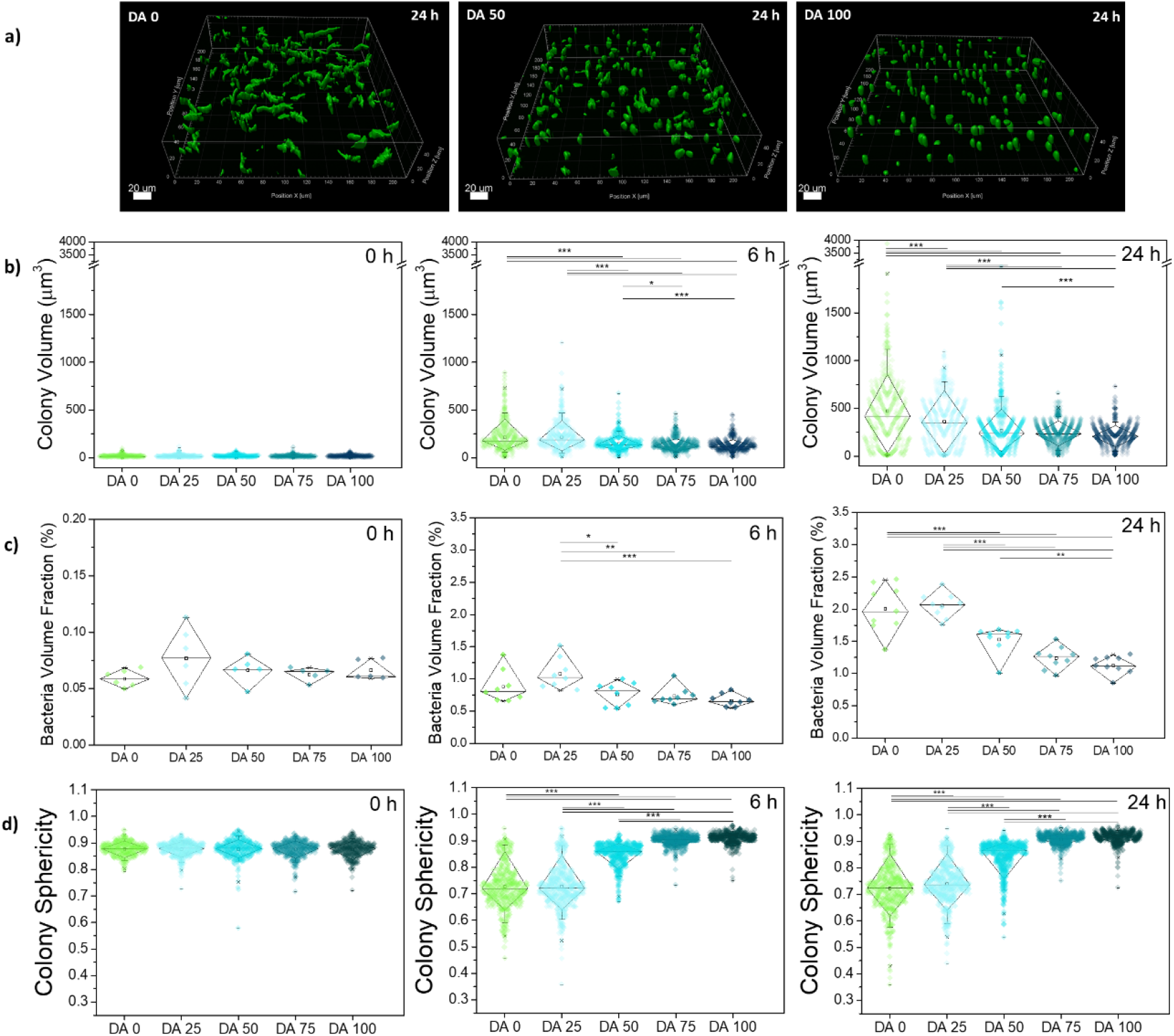
Quantification of bacterial growth in 3D within DA 0-100 matrices formed in microwells. **a)** Exemplary volumetric scan of iLOV producing bacterial colonies inside the hydrogels after 24 h. This visualization is presented using surface masks made with the IMARIS software (scale = 20 μm), **b)** Quantified colony volume within DA 0-100 hydrogels at 0, 6 and 24 h timepoints; **c)** Volume fraction of bacterial colonies within DA 0-100 hydrogels at 0, 6 and 24 h timepoints. **d)** Sphericity values for individual colonies within DA 0-100 hydrogels at 0, 6 and 24 h timepoints. (Diamond plots indicate median, 10 and 90 percentile values, whiskers indicate 5 and 95 percentile values, 338 ≥ Number of colonies ≥ 667 from 3 individual experiments, * p < 0.05, ** p < 0.01, *** p < 0.001)

The sphericity of the colonies increased with the degree of chemical cross-linking (Figure 3d, **Figure S10**), in agreement with the trends previously observed in the in-plane measurements and aspect ratio (Figure 2, Figure S5c, S5f). The distribution of bacterial volume fraction and colony sphericity was narrower in hydrogels with higher degree of covalent cross-linking (Figure 3c, d), also in line with the in-plane measurements (Figure 2). Additional experiments with iLOV producing bacteria in microchannels (**Figure S11**) showed similar trends, indicating that the control mechanism of bacterial colony growth and morphologies imparted by the chemical crosslinks (i.e. by the mechanical constraint) is independent of metabolic variations.

We then explored whether the metabolic activity of the encapsulated bacteria was also influenced by the mechanical constraint of the hydrogel network. It is important to note that induced protein overexpression competes with metabolic resources needed for bacterial growth. Strategies to externally regulate these two processes are of utmost relevance in biotechnological production chains to maximize production yield.^[34-36]^ For our analysis we used a red fluorescent protein (RFP)-producing strain which can be induced by light to drive overexpression of the protein.^[12]^ Since our previous results indicated bacterial growth within the first few hours (Figure 2), induction was initiated right after encapsulation. RFP production was detected after 4-5 hours and it increased linearly with time in all hydrogels up to 11-12 hours, at which time the signal saturated (**Figure 4b**). Interestingly, the extent of protein production, quantified at 10 h, was higher in hydrogels with intermediate degrees of chemical cross-linking (DA25 and DA50) (Figure 4c). To check whether this behavior was driven by a confinement-dependent balance between metabolic activity and growth, a complementary experiment was performed in which encapsulated bacteria were first allowed to grow for 6 h (rapid growth phase) in the dark (no induction) after which protein production was induced by light. Under these conditions, the extent of RFP production did not vary appreciably across the different gel compositions (**Figure S12**), even though the gels with higher cross-linking degrees would have a lower number of cells. This suggests that the slower growing colonies in hydrogels with higher chemical crosslinking were more efficient in RFP production compared to colonies growing faster in physical hydrogels. Thus, bacterial growth restrictions imposed by the mechanical properties of the Plu/PluDA hydrogel network regulated the metabolic activity of the encapsulated cells.

**Figure 4.**
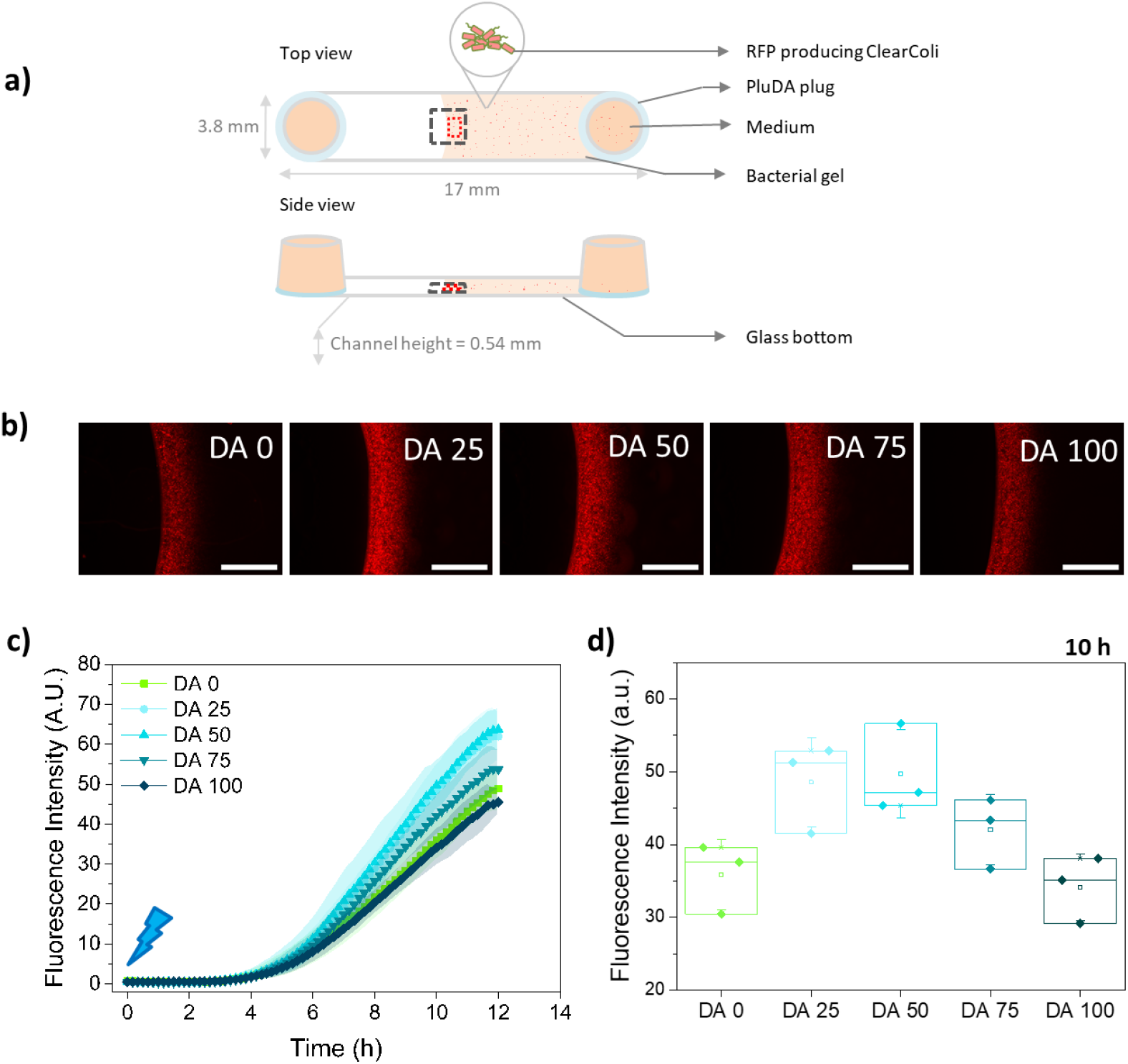
Quantification of protein production by bacteria encapsulated in DA 0-100 hydrogels. **a)** Schematic of the bacterial gels within the microchannels with an open end. The black dotted box (3636 × 2727 um^2^) represents the observed field of view in the microscope, taken at ca. 0.27 mm height, and the red dotted box (200 × 600 um^2^) is the area considered for RFP intensity measurement; **b)** Fluorescence images of RFP producing bacterial gels indicating RFP expressed by the encapsulated bacterial colonies at 10 h (induction i.e. blue light illumination started at 0 h, scale = 1000 μm); **c)** Quantification of fluorescence intensity indicating RFP production in the hydrogels during 12 h (mean ± standard deviation); **d)** RFP production values within different hydrogel compositions at 10 h (N = 3, box represents 25 and 75 percentile values, whiskers indicate standard deviation).

These results show that hydrogel networks combining reversible and permanent crosslinks can be used as external modulators of the growth and metabolic activity of cells in ELMs, although a deeper investigation is required to understand and fully exploit the promising non-monotonic effects observed in our studies.

## 3. Discussion

During biofilm formation, soil homeostasis or invasion of biological tissues, bacteria need to push against their natural surroundings to accommodate space for new cells and grow as colonies. The resulting contact pressures, which are dependent on the mechanical properties of the local microenvironment, have been shown to influence bacteria cell size^[23]^ and to slow down cell growth and delay cell cycle in bacteria studies in synthetic model systems,^[37,38]^ though the principles and mechanisms behind this behavior remain to be investigated. Most studies of bacteria in spatial confinement have been performed in microchannels of different dimensions^[39]^ or in microdroplets.^[40]^ Hydrogel networks are alternative models for such studies. Reported cases have used physical hydrogel networks formed by reversible crosslinks that can rearrange dynamically as bacteria push while growing into larger colonies. The DA 0-100 hydrogels used in our work contain physical and covalent (i.e. permanent) crosslinks at adjustable ratios and yield encapsulating materials with tunable viscoelasticity within a range that influences the embedded *E. coli*. In these hydrogels, bacteria form compact, viable and metabolically active colonies of smaller size and at slower rate when the ratio of permanent crosslinks increases.

Rod-shaped bacteria exert longitudinal forces during growth, contributed by its turgor pressure (30 – 300 kPa for *E. coli*) and limited by the rate of cell wall synthesis.^[21]^ The stress applied to a surrounding hydrogel has been estimated around ∼10 kPa for *E. coli* at their ends.^[41]^ This stress is enough to deform physically crosslinked hydrogels, where bacteria form large colonies and outgrow if nutrients are available.^[10]^ Plu/PluDA hydrogels with PluDA fraction >25 % restrict bacterial growth and colony size, independent of nutrient availability, in a way that seems to correlate with their higher mechanical resistance to deformation. This allows biocontainment of the bacterial population inside the hydrogel matrix, a highly desired feature in engineered living materials and related devices.^[42]^ 25-30 wt% Plu based hydrogels have been used to encapsulate bacteria (*Escherichia coli*)^[29]^ and yeast (*Saccharomyces cerevisiae*)^[16,17]^ in ELMs, and maintain bacteria viability for months.^[16]^ Such Plu/PluDA based hydrogels are stable in vitro for days^[23,43]^ to years ^[16]^ and in vivo for weeks^[44]^ to months.^[45]^ The mechanisms behind bacterial regulation by mechanical confinement remain elusive, but the Plu/PluDA compositions studied in this article seem to be appropriate models to investigate them.

Beyond cellular growth, a recent report demonstrated that photosynthesis in cyanobacteria can also be regulated by mechanical confinement,^[46]^ suggesting that contact pressure could be used as external factor to regulate metabolic processes in bacteria. The viscoelasticity-dependent behavior of mammalian cells in dynamic matrices is well documented and has played a vital role in advancing the fields of 3D cell culture and tissue regeneration.^[27]^ The correlation between bacterial behavior and matrix mechanics, summarized in **Figure 5**, demonstrates that bacterial behavior in confinement also follows mechanical rules of the microenvironment, though sensitivity might occur within a different range of viscoelasticity compared to mammalian cells.

**Figure 5.**
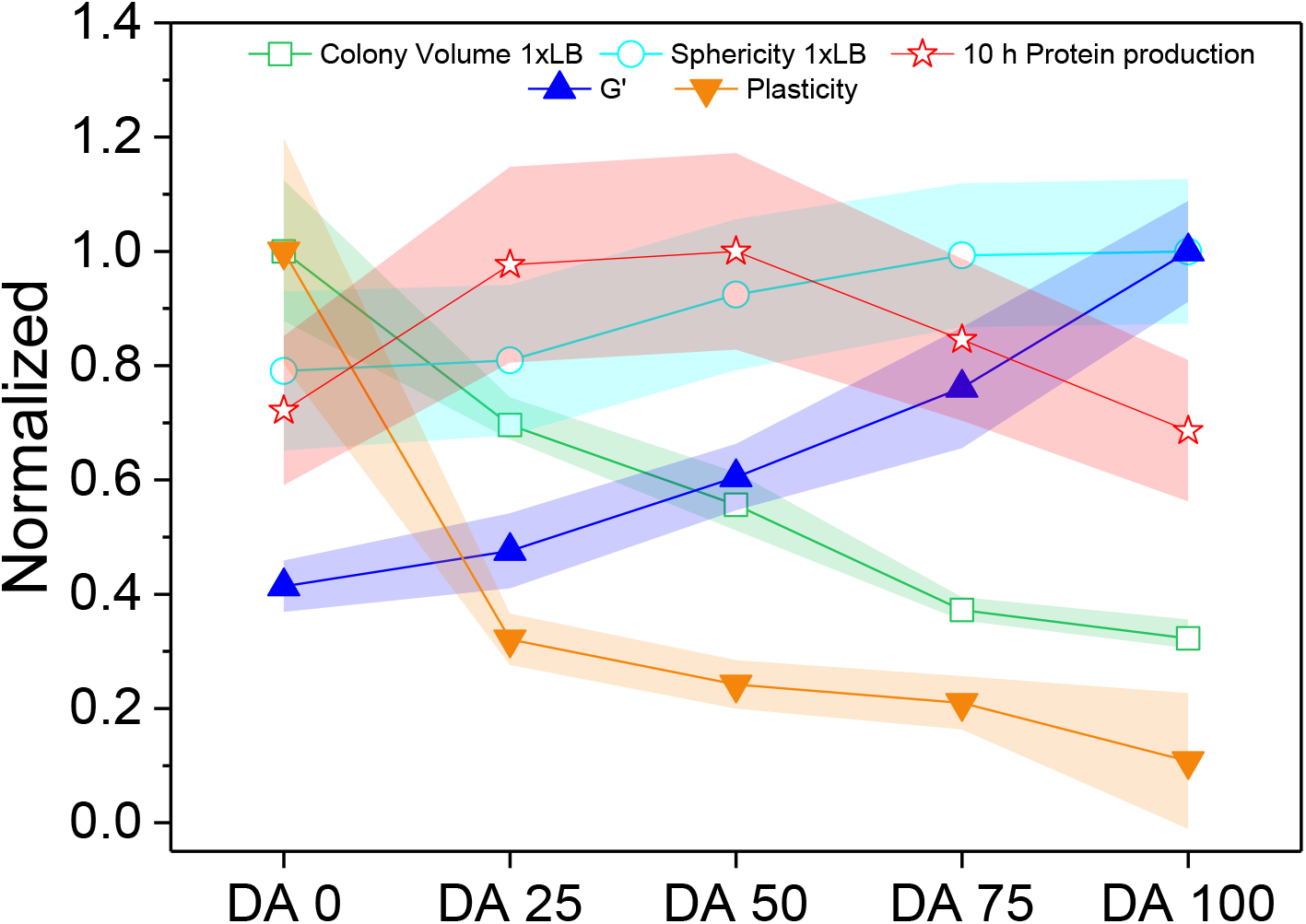
Comparative representation of normalized material’s properties and bacterial response as a function of Plu/PluDA matrix composition. Normalized values of shear storage modulus and degree of plasticity (from Figure 1), 95^th^ percentile volumes and sphericity of colonies after 24 h (from Figure 3), and value protein production after 10 h (from Figure 4). The values were normalized with respect to the highest value in each category (Original values plotted in **Figure S13**). The symbols and shaded regions of the normalized colony volume data indicate 95 ± 2 percentile value range, for all other data the symbols and shaded regions indicate mean ± SD.

## 4. Conclusion

This study comprehensively explores the influences of matrix mechanical properties on the growth and protein production behavior of bacteria encapsulated within viscoeslatically tunable hydrogel networks. We identified behavioral changes occurring solely due to mechanical restrictions imposed by the matrix and not due to nutrient or gas diffusion effects. Our results indicate that bacterial functions can be regulated by careful design of the encapsulating matrix and demonstrates that understanding such behavioral changes in model matrices can enable the optimization of ELM matrix designs to maximize functionality of the embedded organisms. Apart from ELMs, our Pluronic-based mechanically tunable hydrogel platform can also be a powerful tool for gaining fundamental insights into the influence of constrictive forces on bacteria in confined environments (e.g. biofilms).

## 5. Experimental Section/Methods

### Preparation of precursors solutions

Pluronic diacrylate (PluDA) was synthesized by reaction of Pluronic F127 (Plu) with acryloyl chloride in the presence of triethylamine according to a reported protocol.^[31]^ 30% (w/v) Plu and PluDA stock solutions in milliQ water (for rheological measurements) or LB medium (for bacterial growth measurements) containing Irgacure 2959 photoinitiator at 0.2% w/v were prepared and stored at 4 °C and allowing them to form the gel at room temperature for 10 minutes.^[47]^ The composition of DA0 and DA100 hydrogels corresponded to that of the stock solutions of Plu and PluDA, respectively. DA 25, DA 50 and DA 75 hydrogels were prepared by mixing of the stock solutions in ratios as shown in **Table 1**. For the photoinitiated crosslinking of the hydrogels, these were exposed to UV light (365 nm, 6 mW/cm^2^) using a OmniCure Series 1500 lamp during 60 s (Milli Q water solutions) or 120 s (LB medium solutions).

### Rheological studies

The rheological properties were measured using a rotational rheometer (DHR3, TA Instruments) using a parallel plate geometry. A 20mm Peltier plate/ UV transparent plate was used as bottom plate and a smooth stainless steel 12 mm disk was used as top plate. The rheometer was equipped with a UV Source (OmniCure, Series 1500, 365 nm, 6 mW/cm^2^) for illumination of the hydrogel samples between the rheometer plates. Experiments were performed at room temperature unless otherwise mentioned. To avoid drying of the sample by evaporation during testing, a solvent trap was used, and the sample was sealed with silicone oil. A volume of precursor solution of 35 µl and a gap between plates of 300 µm was used for the experiments. All tests were initiated 10 minutes after loading the sample on the rheometer. Strain sweeps were conducted from 0.001 to 1000% at a frequency of 1 Hz. Frequency sweeps were conducted from 0.01 to 100 Hz at a constant strain of 0.1% with a controlled temperature of 23 °C. From these experiments, the linear viscoelastic region was identified (Figure S1a, S1b and S1c). Temperature sweep experiments were carried out from 4 to 40 °C at a 5 °C/min ramp rate, 1 Hz frequency and 0.1% strain value (found to be in the linear viscoelastic range). These experiments were used to characterize the gelation temperature of the solutions (Figure S1d). The gelation point (defined as crossover point of G’ and G’’) in the temperature sweep experiment was 15.5 °C for 30% DA 0 solutions and 14 °C for DA 100. The shear modulus of DA 0 and DA 100 before crosslinking was similar (Figure S1). Time sweep experiments to monitor the changes in the mechanical properties of the hydrogels during crosslinking were performed. For this purpose, samples were irradiated at 365 nm (6 mW/cm^2^) during 60 s at room temperature, and the changes in the shear moduli were recorded for a total of 7 minutes (Figure S2). All time sweep experiments were performed at least in triplicate.

### Creep recovery experiments

For creep-recovery experiments a shear stress of 100 Pa was applied for 3 minutes to the hydrogel sample in the rheometer. The shear strain was monitored for during this time (creep phase), and for further 3 min after removal of the shear stress (recovery phase). The maximum strain during the load phase and the residual strain after recovery were obtained from the rheology curves and represented as function of hydrogel composition. The maximum strain was defined as the highest value of strain achieved during stress application. The residual strain was defined as the steady-state strain after 3 minutes recovery. The ratio between the irreversible strain and the peak strain was defined as the degree of plasticity.^[48]^

### Bacteria cultures

An endotoxin free strain of *E. coli* (ClearColi^(R)^ BL21(DE3), BioCat) ^[12]^ was used for the bacterial growth studies. It was transformed with the plasmid pUC19 to enable Ampicillin resistance and minimize the risk of contamination in the culture. Bacterial cultures were grown for 16h at 35°C, 180 rpm in LB Miller medium supplemented with 50 µg/mL of Ampicillin to an optical density at 600 nm wavelength (OD600) value between 0.5 – 1. For the fluorescence based experiments, we used a previously optogenetically engineered strain of ClearColi that produces red fluorescence protein (RFP) when illuminated with blue light,^[12]^ and a ClearColi strain that constitutively produces iLOV protein. The iLOVf gene insert was amplified from the pET28-iLOVf plasmid (Addgene #63723) and assembled into the pLp-3050sNuc vector^[49]^ (Addgene #122030) using Gibson Assembly. The iLOVf gene is under the constitutive expression of the P48 promoter.^[50]^ The construct was sequence verified and subsequently transformed in the Clearcoli BL21 DE3 strain and maintained with the additional supplementation of 200 µg/mL Erythromycin. Since these bacteria grew slightly slower than the pUC19 harboring variants (Figure S8), they were initially cultured for 16h at 35°C, 220 rpm in LB Miller medium until an optical density at 600 nm wavelength (OD600) value of 0.2 – 1 was reached. The RFP producing strain harbors the plasmid pDawn-RFP that encodes blue-light activatable gene expression and provides kanamycin resistance. The red fluorescence signal was used to image bacterial growth inside the hydrogels and to quantify protein production in hydrogel-encapsulated bacterial populations. The bacteria were cultured for 16h at 35°C, 220 rpm in LB Miller medium supplemented with 50 µg/mL of Kanamycin to an optical density at 600 nm wavelength (OD600) value between 0.5 – 1.5. All procedures with the light-inducible bacterial strain were performed either in the dark or under a laminar hood with an orange film that cuts off blue light.

### Bacterial encapsulation in hydrogels

Hydrogels with 30% (w/v) polymer concentration were prepared by mixing stock solutions of Plu and PluDA precursors in bacterial medium with bacterial suspensions (at OD_600_ of 0.5 ∼ 4 × 10^7^ cells/mL) at 9/1 (v/v) ratio to achieve a final OD600 of 0.05 within the gels. This concentration allowed individual bacteria to be homogeneously dispersed in the hydrogels and most to grow into individual bacterial colonies that did not coalescence during 24 h. This allowed quantification of individual colony dimensions and morphologies throughout the hydrogel. The solutions were stored in ice before and after mixing. At this temperature Pluronic solutions are liquid and can be homogenously mixed with the bacteria by pipetting at the start of the experiment.^[45]^ Hydrogels were produced in two different formats: (i) Inside channels using Ibidi μ-Slides VI 0.4 (17 × 3.8 × 0.4 mm^3^) or Ibidi µ-Slide VI 0.5 Glass Bottom (17 × 3.8 × 0.54 mm^3^) consisting of 6 parallel microfluidic channels and (ii) inside ibidi μ-Slide angiogenesis micro-wells with polymer coverslip bottom (Figure S3). For the in-plane colony length extension analysis (Figures 2a), Ibidi μ-Slides VI 0.4 were used. 30 µL of this pluronic/bacteria suspension was pipetted into the microchannel, placed on ice, filling the entire channel. For the high magnification confocal microscopy experiments to determine the colony volume (Figure S11), 30 µL of this pluronic/iLOV-producing bacteria suspension was pipetted into the Ibidi µ-Slide VI 0.5 Glass Bottom microchannel. Experiments in Figure 3a were done with Pluronic/iLOV-producing bacteria suspensions (10 µL) pipetted into the micro-well of the μ-Slide angiogenesis dish. Measurements were done near the center of the wells and 50 µm from the bottom. Ibidi µ-Slides VI 0.5 with glass bottom were used for the experiments with pluronic/RFP-producing bacteria suspension (Figure 4). 20 µL suspension was pipetted into the microchannel and filled up to half the length of the channel, resulting in a sharp hydrogel-air interface (Figure 4), to enable oxygen availability required for folding of the RFP protein.^[12]^ RFP production was measured 50 µm from the edge of the gel-air interface using a 200 × 600 µm^2^ area as the region of interest. The reservoir at both the ends of the channel were plugged with 20 µL of DA 100 gel and 50 µL of medium to avoid drying of the hydrogel in the channel for all the experiments with channels. The microculture was kept at room temperature for 10 min for physically cross-linked gelation to occur and exposed for 120 s to a UV Lamp inside Alpha Innotech FluorChem Q system (Biozym, Oldendorf, Germany) (6 mW/cm^2^) which was the illumination step used to initiate the photopolymerization of the acrylate groups and covalent crosslinking of the hydrogels. The bacterial gels in μ-Slide angiogenesis micro-wells were topped with 20 µL of silicone oil (350 cSt, Sigma-Aldrich) to prevent drying of the hydrogel during the experiment.

### Imaging and quantification of in-plane length extension of colonies inside the channels

Brightfield microscopy analysis was performed using Nikon Ti-Ecllipse (Nikon Instruments Europe B.V., Germany) microscope with 20× S Plan Fluor objective with a numerical aperture of 0.45 and a working distance of 8 mm, Sola SE 365 II (Lumencor Inc., Beaverton, USA) solid state illumination device and an Andor Clara (Andor Technology Ltd, Belfast, United Kingdom) CCD camera for detection. Ibidi μ-Slides VI loaded with the bacterial hydrogels were incubated at 37 °C and 5 % CO_2_ using an Okolab (Okolab SRL, Pozzuoli, Italy) incubation chamber coupled to the microscope. Imaging locations were selected near the middle of the channel length and about halfway between the bottom and top of the channel. Such a position was chosen to minimize the possibility of variations in material properties (swelling, dissolution) or nutrient variability that might occur due to diffusion of medium at the reservoirs. Changes in material properties of the gel near the reservoirs was inferred from movement of the bacterial gel in and out of the imaging field during the experiment. Time-lapse imaging was performed from 30 min to 1 h after introducing the bacterial gels in the channel slides and subsequently in the incubation chamber to ensure that the samples reach 37 °C. Time-lapse imaging was done with an interval of 15 minutes for 12 h. Each type of experiment was performed in triplicates.

### Quantification of colony growth

The pluronic/iLOV-producing bacteria samples in the ibidi μ-Slide angiogenesis micro-wells were imaged by Zeiss LSM 880 confocal laser scanning microscopy (CLSM) at 0 h, 3 h, 6 h and 24 h timepoints. Two-photon laser was used for imaging the iLOV protein producing bacterial colonies.^[51]^ The exposure conditions were optimized for minimizing cell photodamage using the objective LD C-Apochromat 40x/1.1 W Korr M27, detection wavelength 499-624 nm, laser wavelength of 880 nm and power of 5 %. Z-stacks of 50.102 µm were taken in steps of 0.65 µm. Images of a size of (xy) 212.5 × 212.5 µm were acquired with a resolution 512 × 512 pixels, two-fold line averaging, and using constant values for laser power (5 %) and pixel dwell time of 2.06 µs. The digital gain value was 800 for 0 h, 3 h and 6 h measurements and adjusted to 730 for 24 h measurements, as some pixels reached oversaturation. Imaris software (Version 9.0, Bitplane, Zurich, Switzerland) was used to process CLSM image z-stacks to create three-dimensional images using the Imaris surface tool. The surfaces were generated with smooth function set to 0.5 μm, the background threshold to 10 μm and minimum surface voxels limit to 10. The sphericity was quantified as the ratio between the volumes of the colony to its surface area (Imaris surface tool). Volume fraction of the bacterial colonies covering the hydrogel samples was calculated as the ratio of sum of all colony volumes and the volume of the observed hydrogel field of view. For calculation of single colony volumes, the colonies touching the edges of the field of view and those which coalesced together were avoided. Colonies below the volumes of 5 μm^3^ (which was approximately the volume of single bacterium) were neglected.

### Protein production in the hydrogels

Ibidi µ-Slide VI 0.5 Glass Bottom microfluidic channels loaded with bacterial hydrogels were used in these experiments. The culture was kept in static conditions at 37 °C using an incubation chamber coupled to a BZ-X800 (Keyence, Osaka, Japan) microscope. For fluorescence imaging to detect the RFP production, filter channel TRITC (BZ-X Filter OP-87764, excitation 545/25, emission 605/70) was used. RFP production was activated, at 0 h/6 h, by illuminating the channels with blue light (BZ-X Filter GFP OP-87763, excitation 470/40, emission 525/50) pulses of 500 milliseconds every 10 min for 18 hours using the 4X objective (Keyence Plan Apochromat, numerical aperture 0.20, working distance 20 mm) of the microscope. The light intensity and exposure settings were optimized for detecting early time-point generation of fluorescence and following increase in intensity for several hours.

Image processing and analyses were performed using Fiji edition of ImageJ (ImageJ Java 1.8.0). Quantification of the fluorescence intensities was done by determining the mean grey value, at a height of ca. 0.27 mm (mid-point of channel thickness) and 50µm away from the edge (as larger colonies were observed near edge, owing to possible differences in the mechanical properties at the interface with air ^[20,21]^) within a 200 ×600 µm^2^ area of the gel and subtracting the mean grey value of the background.

### Statistical analyses

One-way analysis of variance (ANOVA) with post hoc Tukey HSD test was performed with results involving more than three data points. Differences were considered statistically significant at * p < 0.05, ** p < 0.01, *** p < 0.001. Analyses were performed with Origin Pro 9.1 software.

## Supporting Information

Supporting Information is available from the Wiley Online Library or from the author.

## Acknowledgements

We acknowledge support from the Deutsche Forschungsgemeinschaft’s Collaborative Research Centre, SFB 1027 and the Leibniz Science Campus on Living Therapeutic Materials, LifeMat.

pET28-iLOVf plasmid was a gift from Prof. Brian Smith, Addgene #63723. pLp-3050sNuc vector was a gift from Prof. Geir Mathiesen, Addgene #122030 We also thank Dr. Mitchell Han for his valuable insights in image analysis and Sourik Dey for iLOV plasmid transformation.

Received: ((will be filled in by the editorial staff))

Revised: ((will be filled in by the editorial staff))

Published online: ((will be filled in by the editorial staff))

## Supporting Information

### Synthesis of Pluronic diacrylate (Plu-DA)

Dry Plu-DA (20 g, 1.59 mmol) and triethylamine (0.55 ml, 3.98 mmol) dissolved in 200 ml of dichloromethane were charged in a 500-ml round-bottomed flask. Acryloyl chloride (0.25 ml, 6.36 mmol) was added dropwise. The reaction mixture was stirred at 4 °C for 12 h and then at room temperature for 12 h. A white precipitate formed and was filtered. The solution was concentrated and filtered again. Precipitation in diethylether and drying in vacuum for 1 day rendered 18 g of Pluronic-DA. The chemical structure of Plu-DA and the extent of acrylation were determined by ^1^H-NMR. The acrylation degree was determined by the ratio of the terminal acrylic protons (5.98-6.52 ppm) to the methylene groups of the backbone (1.05-1.28 ppm). Plu-DA with substitution degree of 70% was obtained.

**Figure S1.**
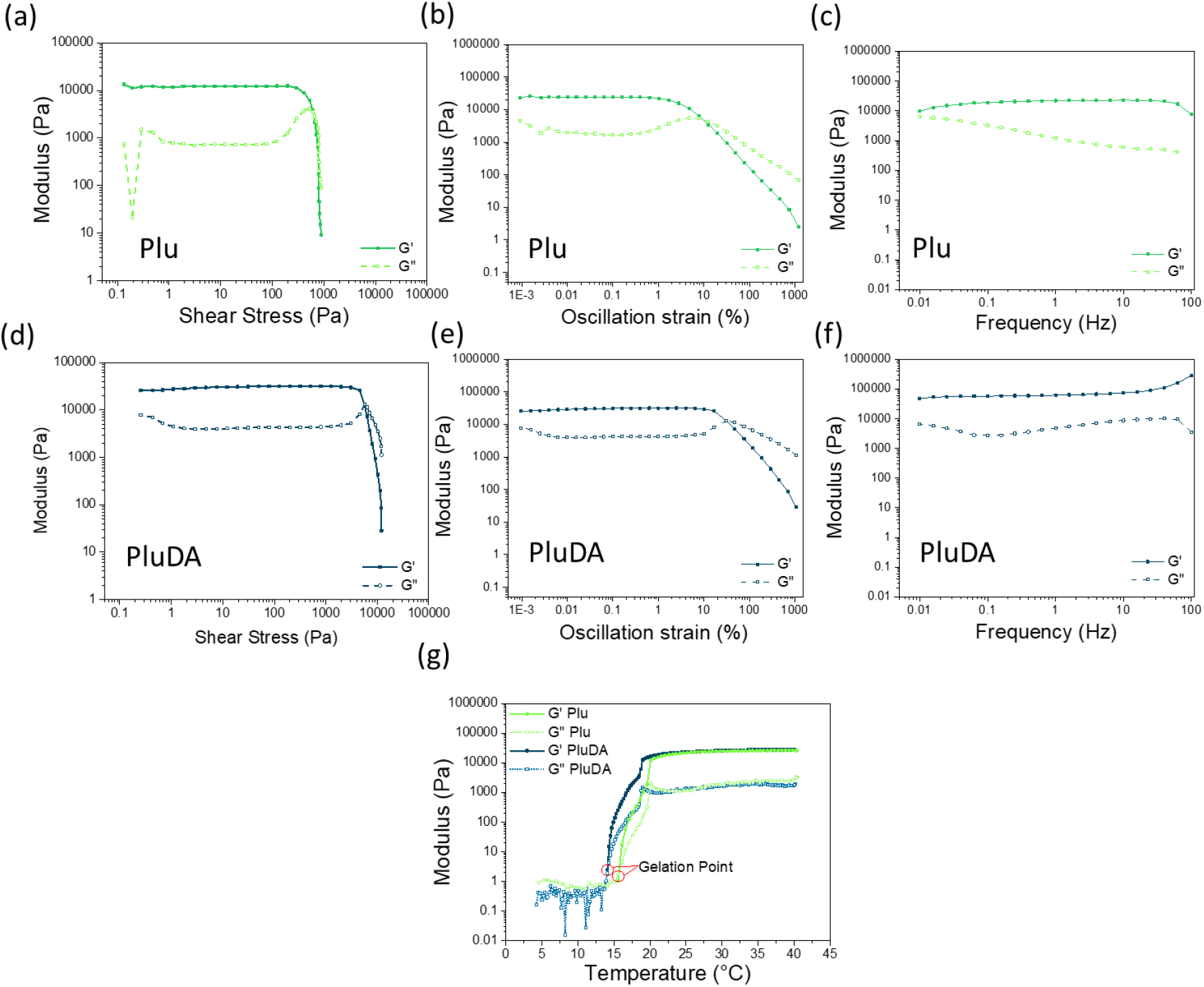
Rheological measurements of Plu and PluDA hydrogels at 30% w/v polymer concentration. **a)** Strain sweep experiment to identify the yield stress value of Plu hydrogel, **b)** Strain sweep and **c)** frequency sweep experiments to determine the linear viscoelasticity region of Plu hydrogel. **d)** Strain sweep experiment to identify the yield stress value of PluDA hydrogel, **e)** Strain sweep and **f)** frequency sweep rheograms to determine the linear viscoelasticity region of PluDA hydrogel. **a-f**: All measurements were performed at room temperature. **g)** Temperature sweep measurement to determine the gelation point (G’= G’’) of Plu and PluDA

**Figure S2.**
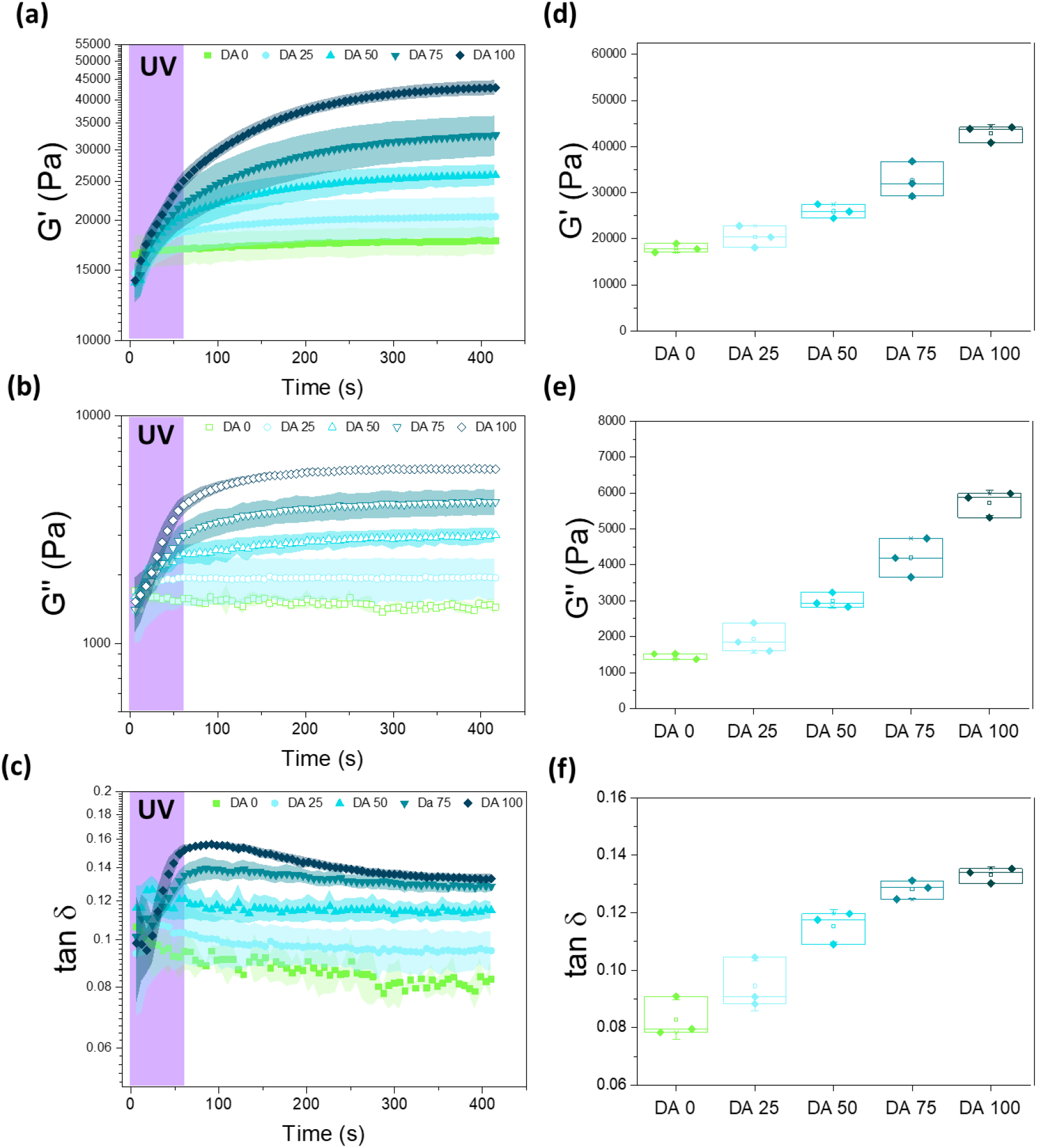
Rheological properties of Plu/PluDA (DA 0-100) hydrogels. **a-c)** Evolution of the **a)** shear storage modulus, **b)** shear loss modulus and **c)** tan δ values after physical gel formation (t=0 s), during photoinitiated crosslinking (t=0-60 s) and afterwards (t=60-400 s). The curves show post-exposure polymerization characteristic of crosslinking processes as a consequence of the hindered mobility of the initiated radicals that slows down the polymerization and termination reactions.^[1]^ **d-e)** Values of **d)** shear storage modulus, **e)** shear loss modulus and **e)** tan δ at t = 420 s from the rheological curves in **a-c**. (N = 3, box represents 25 and 75 percentile values and whiskers indicate standard deviation.)

**Figure S3.**
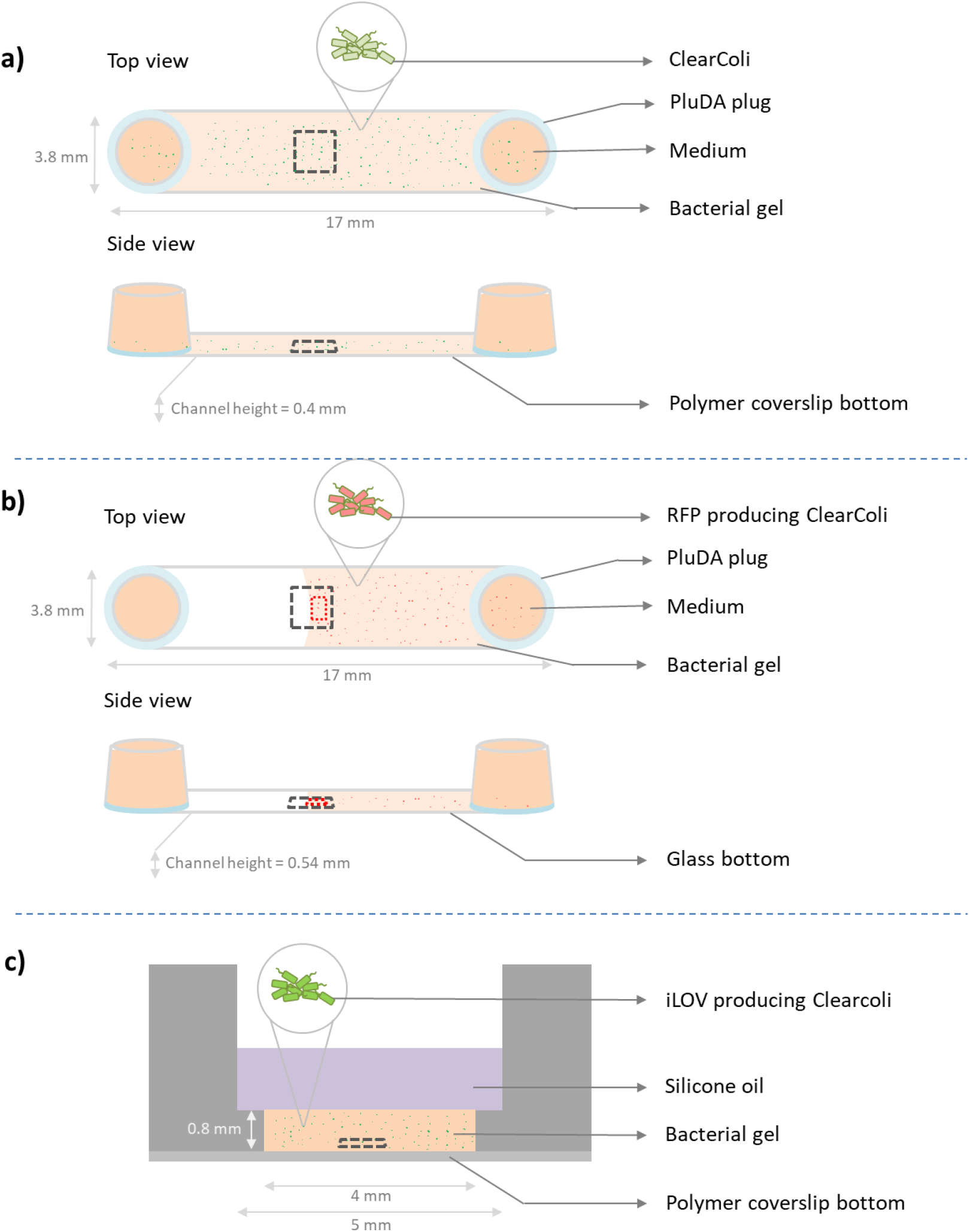
Design of experimental platforms used to quantify bacterial behavior inside the hydrogels. **a)** Microchannels in Ibidi μ-Slides VI 0.4 (17 × 3.8 × 0.4 mm^3^). Volume of bacterial hydrogel used to fill the channel was 30 µL, imaging locations (black dotted box of 449 × 335 um^2^) were selected near the middle of the channel length and nearly 150-250 µm from the bottom of the channel; **b)** Microchannel in Ibidi µ-Slide VI 0.5 Glass Bottom (17 × 3.8 × 0.54 mm^3^). Volume of bacterial hydrogel used to fill the channel was 20 µL, imaging location (black dotted box of 3636 × 2727 µm^2^) representing the observed field of view in the microscope was selected at the hydrogel-air interface at a height of ca. 0.27 mm (mid-point of channel thickness) and quantification of the fluorescence intensities was done 50 µm away from the edge within a 200 ×600 µm^2^ area (red dotted box); **c)** Microwell in Ibidi μ-Slide angiogenesis. Volume of bacterial hydrogel used to fill the well was 10 µL, measurement at 50 µm from the bottom (black dotted box of 212.5 × 212.5 × 50.102 µm^3^). **a-c**) The grey square indicates the region of interest (ROI) of the hydrogel used for imaging, and the red square indicates the region used for quantification of RFP intensity.

**Figure S4.**
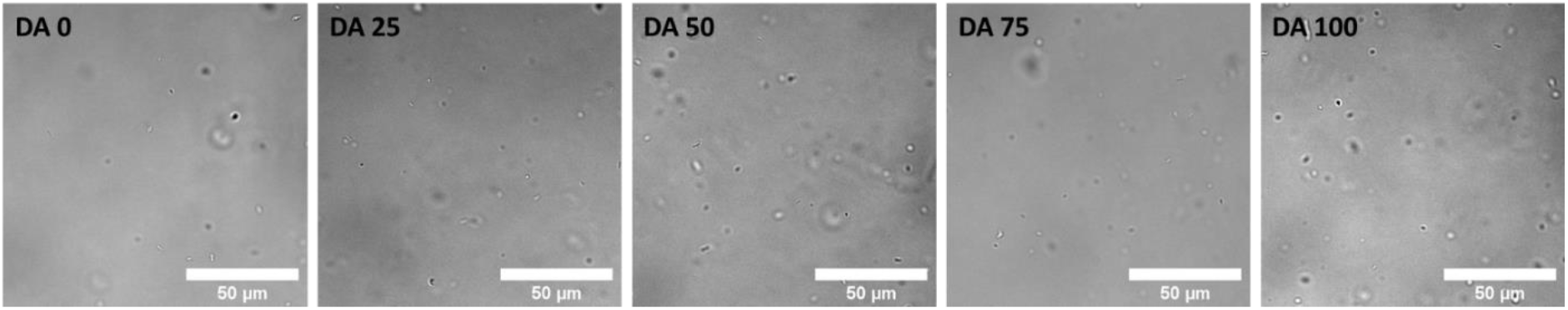
Bright-field images of the bacteria embedded within DA 0, DA 25, DA 50, DA 75 and DA 100 hydrogels with 1x LB concentration at 0 h time point, indicating presence of individual bacterial cells. (scale = 50 μm, done in ibidi μ-Slide angiogenesis microwells)

**Figure S5.**
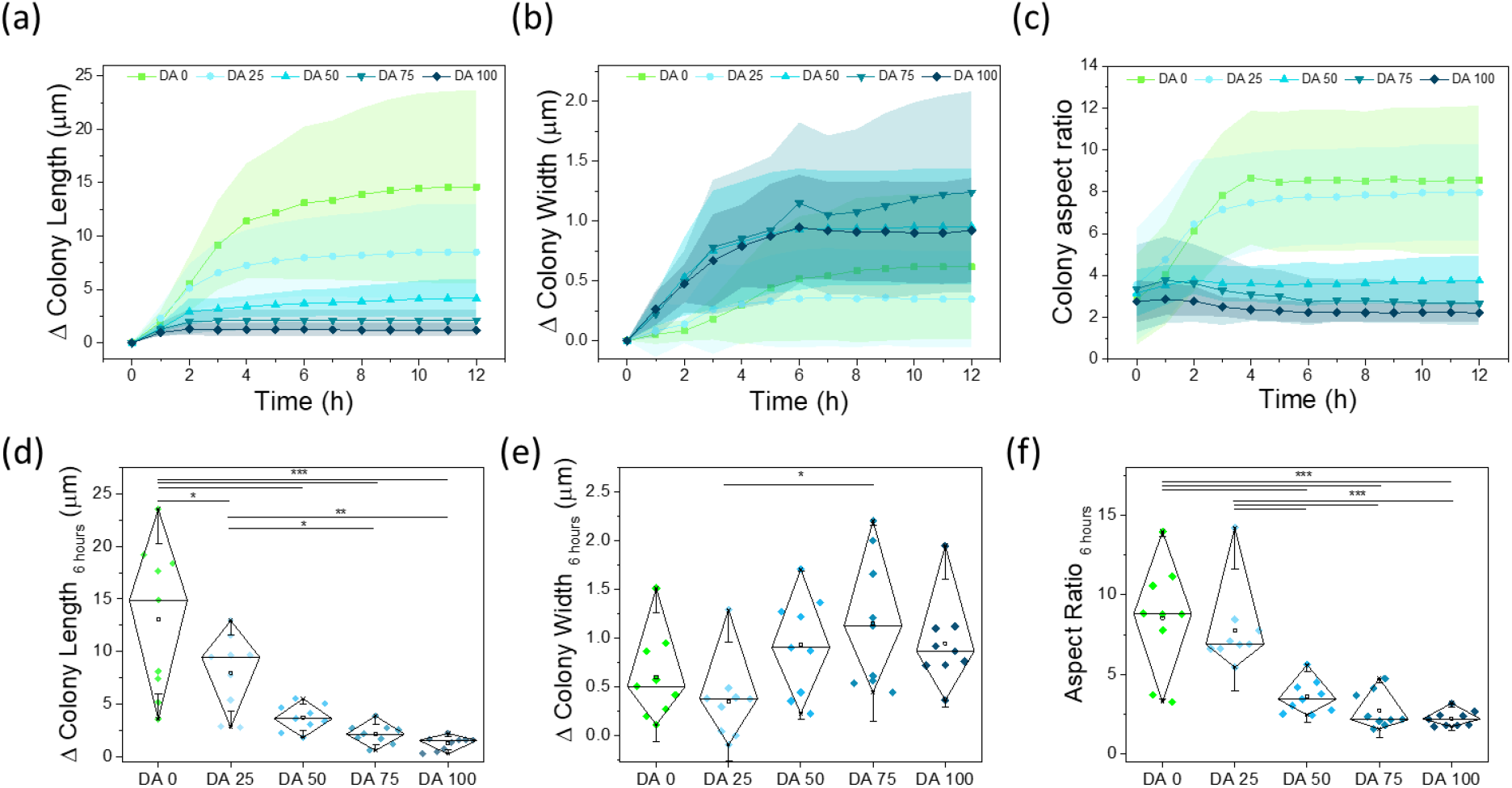
In-plane quantification of dimensional changes of bacterial colonies encapsulated in DA 0-100 hydrogels in microchannels (Figure S3a) with 1x concentration LB medium. **a-c)** Evolution of the length, width and aspect ratio of individual colonies within 12 h of culture (mean ± SD). **d-f)** Comparative values of changes in colony length, width and aspect ratio after 6 hours in the different hydrogels. (Diamond plots indicate 10 and 90 percentile values, whiskers indicate standard deviation values, number of colonies = 9, * p < 0.05, ** p < 0.01, *** p < 0.001)

### Fluorescence recovery after photobleaching (FRAP) in the hydrogels

For all of the FRAP measurements, a fluorescent small molecule erythrosin B was used^[2]^ and observed under a confocal laser-scanning microscope (LSM 880) equipped with an Airyscan detector (Zeiss, Germany). The gels, within the microchannel constructs as shown in Figure S3a, were visualized under an air immersion objective (EC Plan-Neofluar 10x/0.30 Ph1 M27), and images were acquired with a 514 nm laser diode, detection wavelength 519-653 nm, dwell time of 0.42 µs per pixel and 0.1% laser power. For bleaching, a circular region of interest with a diameter of 26 µm was illuminated using a 405-nm laser diode at 30% laser power with a dwell time of 131 µs per pixel. The total acquisition length was 261.9 s, with a frame rate of 2 fps.

**Figure S6.**
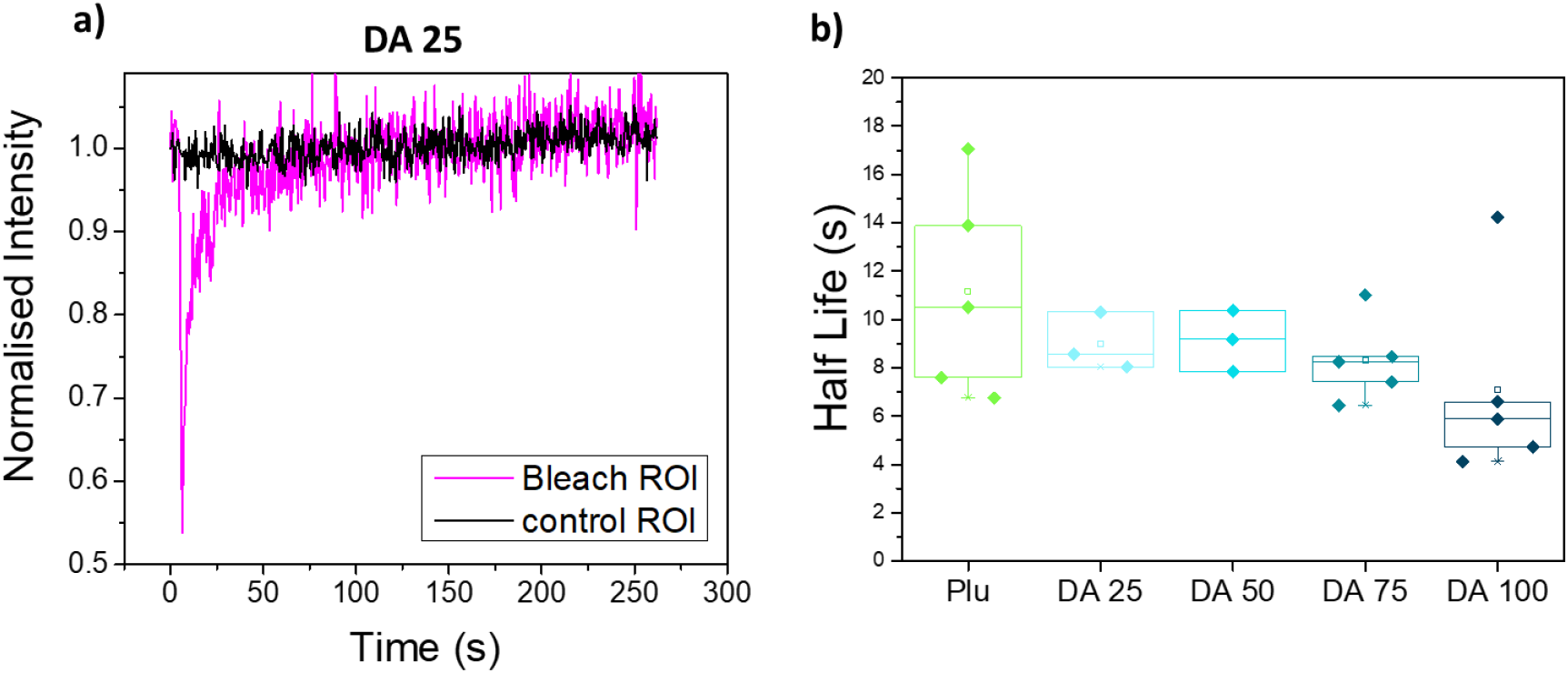
FRAP assay comparison of molecular diffusion rates inside DA 0-100 hydrogels. **a)** Normalized FRAP curve of Erythrosin B dye in DA 25 hydrogel. **b)** Quantified recovery time values, τ ½, of FRAP in the DA 0-100 hydrogels. The obtained τ ½ were close to 10 seconds for all hydrogel compositions with no significant differences. This indicates that the diffusion rate of nutrients within the hydrogels are expected to be similar.

**Figure S7.**
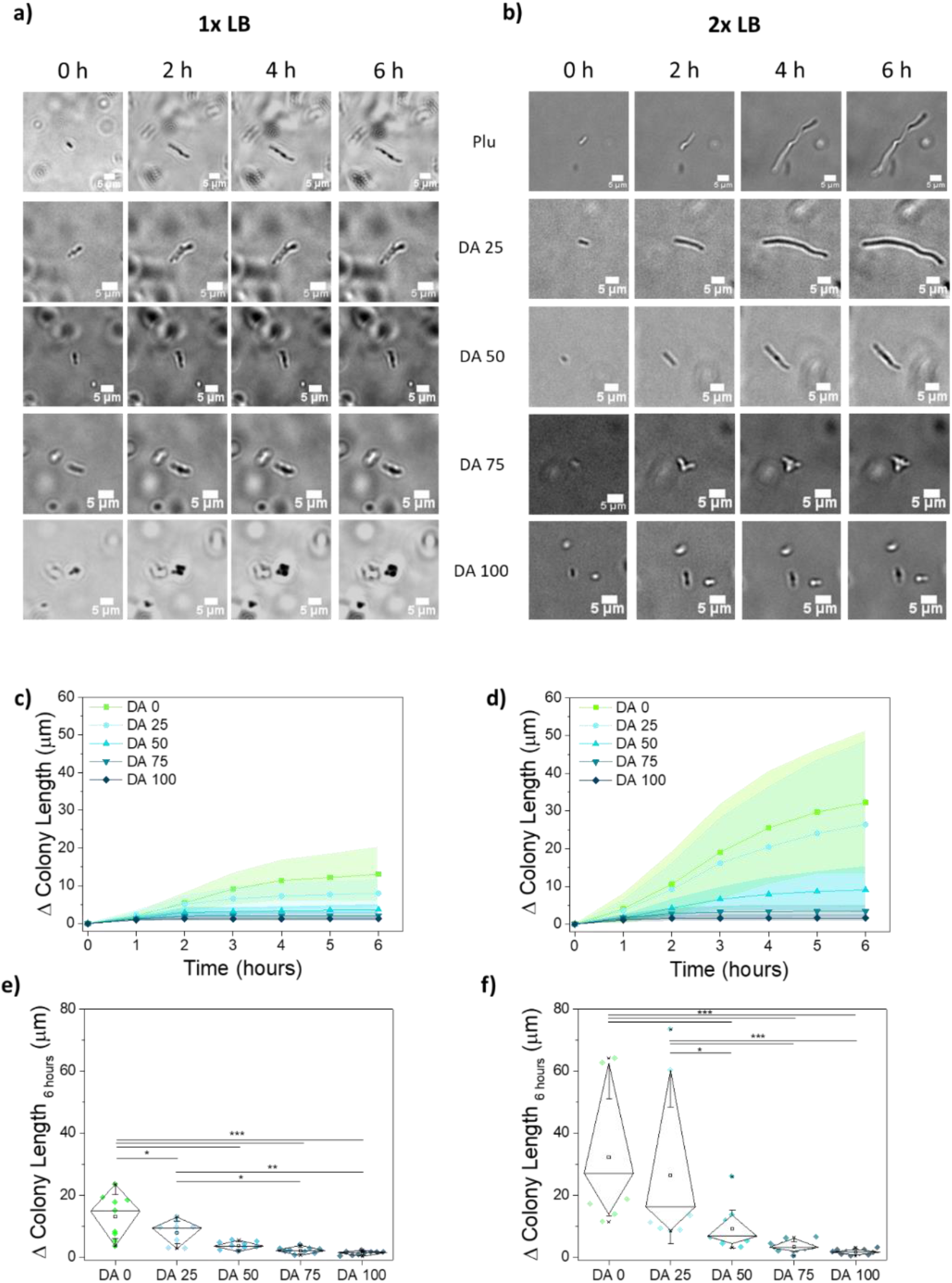
Bacteria colony growth inside hydrogels in microchannels (Figure S3a). **a)** Time lapse bright-field images of bacterial colonies within Da 0-100 hydrogels in 1x LB medium. Scale bar corresponds to 5 μm. **b)** Time lapse bright-field images of bacterial colonies within DA0-100 hydrogels in 2x LB medium. Scale bar corresponds to 5 μm. **c-d)** Quantification of the extension of bacteria colony length within the hydrogels with **c)** 1x and **d)** 2x LB concentration with increasing time; **e**,**f)** Bacterial length extension values at 6 h with **e)** 1x and **f)** 2x LB concentration. (Diamond plots indicate 10 and 90 percentile values, whiskers indicate standard deviation values, 9 ≥ Number of colonies ≥ 13, * p < 0.05, ** p < 0.01, *** p < 0.001)

**Figure S8.**
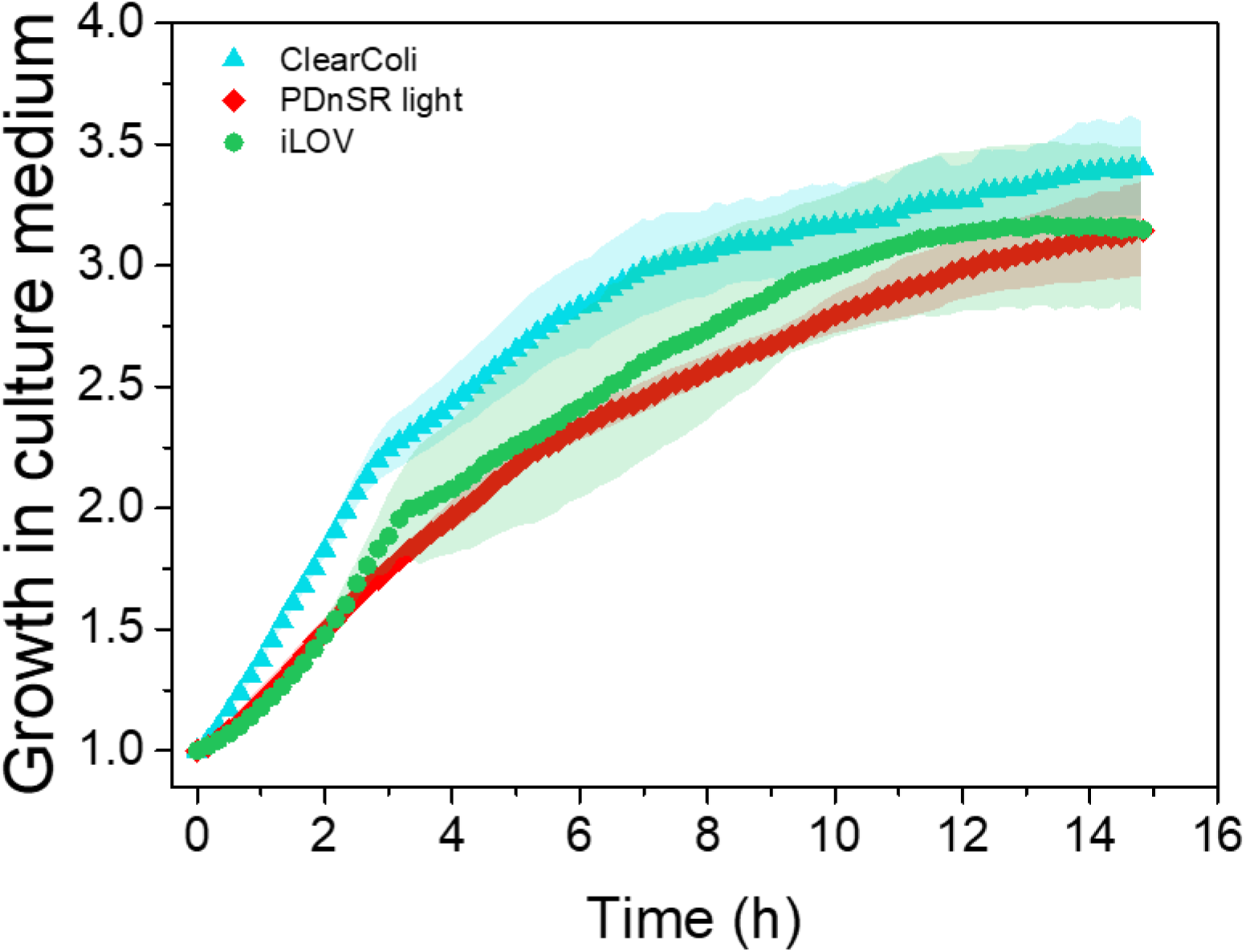
Growth rate of ClearColi, genetically modified RFP producing ClearColi and genetically modified iLOV producing ClearColi bacteria in culture medium determined by OD 600 measurements in a microplate reader (TECAN Infinite 200 Pro) at 37°C with shaking. The y axis indicates the multiplication factor of the growing bacteria population compared to the starting OD 600 of 0.2. (Symbols – mean, shaded regions – SD, N=3)

**Figure S9.**
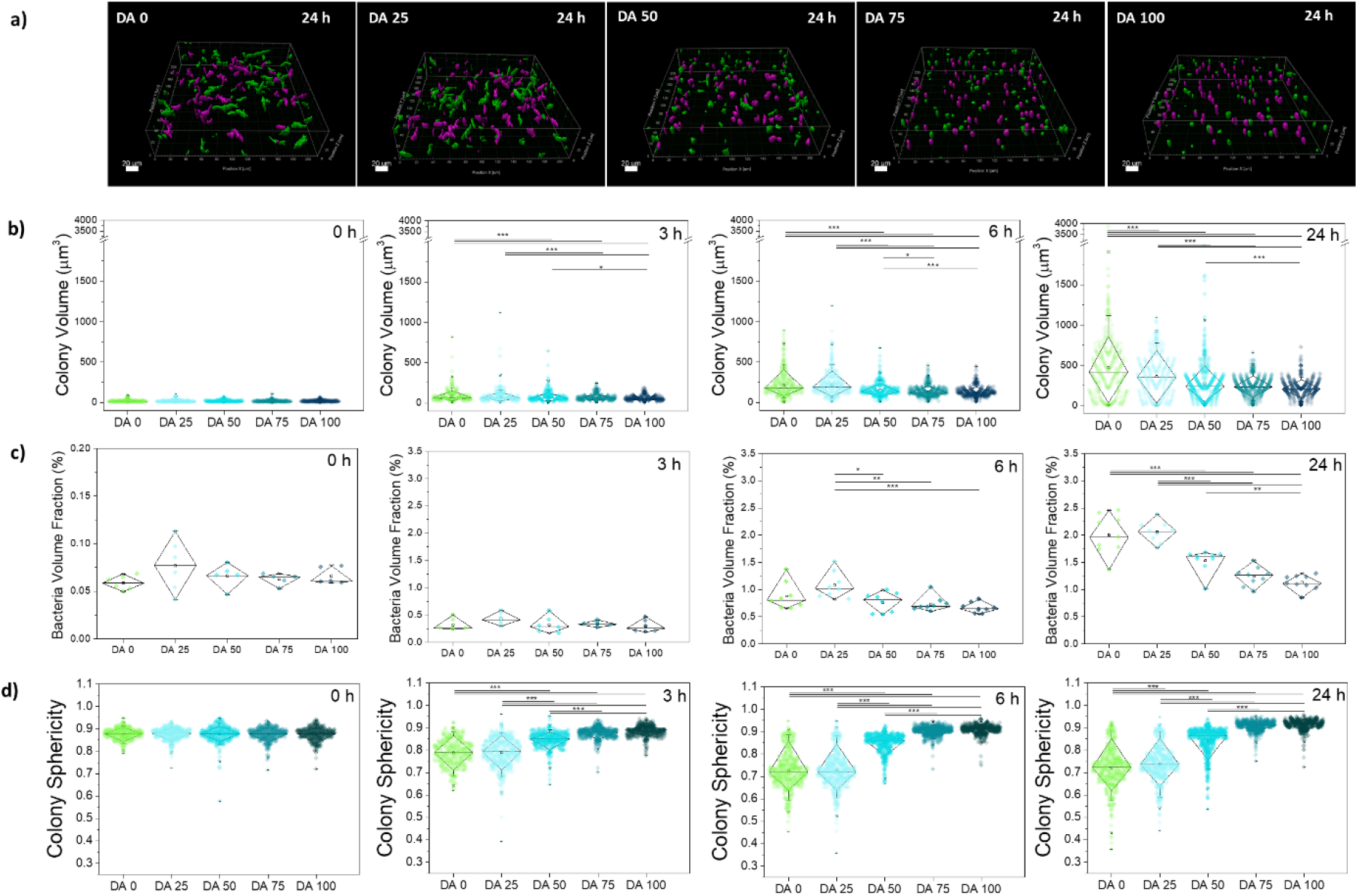
Bacteria colony growth inside hydrogels in microwells (Figure S3c). **a)** Fluorescence image of 212.5 × 212.5 × 50.102 µm^3^ volumes of bacterial hydrogels producing iLOV after 24 h culture. The magenta-colored colonies do not touch the edges of the ROI, while the green colored colonies do. The magenta and green colored colonies were used for the quantification of the bacteria volume fraction **(b)**. Only magenta-colored colonies were considered for the quantification of the volume **(c)** and sphericity **(d)** of individual colonies. For visualization surface masks obtained by IMARIS software were used. The scale bar corresponds to 20 μm. **b)** Volume of individual colonies in DA 0-100 hydrogels quantified at 0, 3, 6 and 24 h timepoints; **c)** Bacteria volume fraction in DA 0-100 hydrogels quantified at 0, 3, 6 and 24 h timepoints; and **d)** Sphericity of bacteria colonies quantified in DA 0-100 hydrogels. (Diamond plots indicate 10 and 90 percentile values, whiskers indicate 5 and 95 percentile values, 338 ≥ Number of colonies ≥ 667, * p < 0.05, ** p < 0.01, *** p < 0.001)

**Figure S10.**
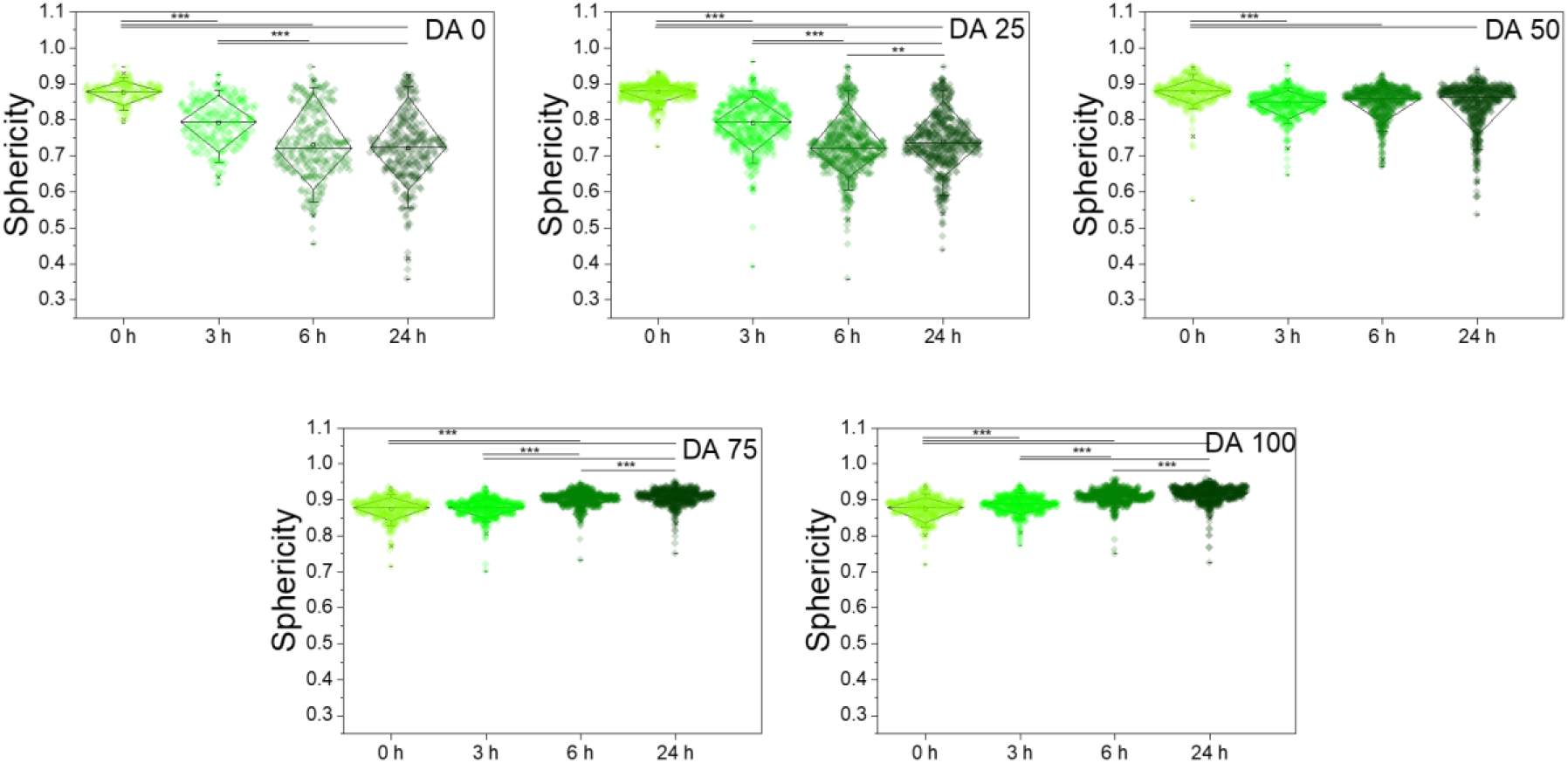
Quantification of the sphericity of iLOV producing ClearColi colonies within DA 0-100 hydrogels in microwells (Figure S3c) as function of culture time. (Diamond plots indicate 10 and 90 percentile values, whiskers indicate 5 and 95 percentile values, 338 ≥ number of colonies ≥ 667, * p < 0.05, ** p < 0.01, *** p < 0.001)

**Figure S11.**
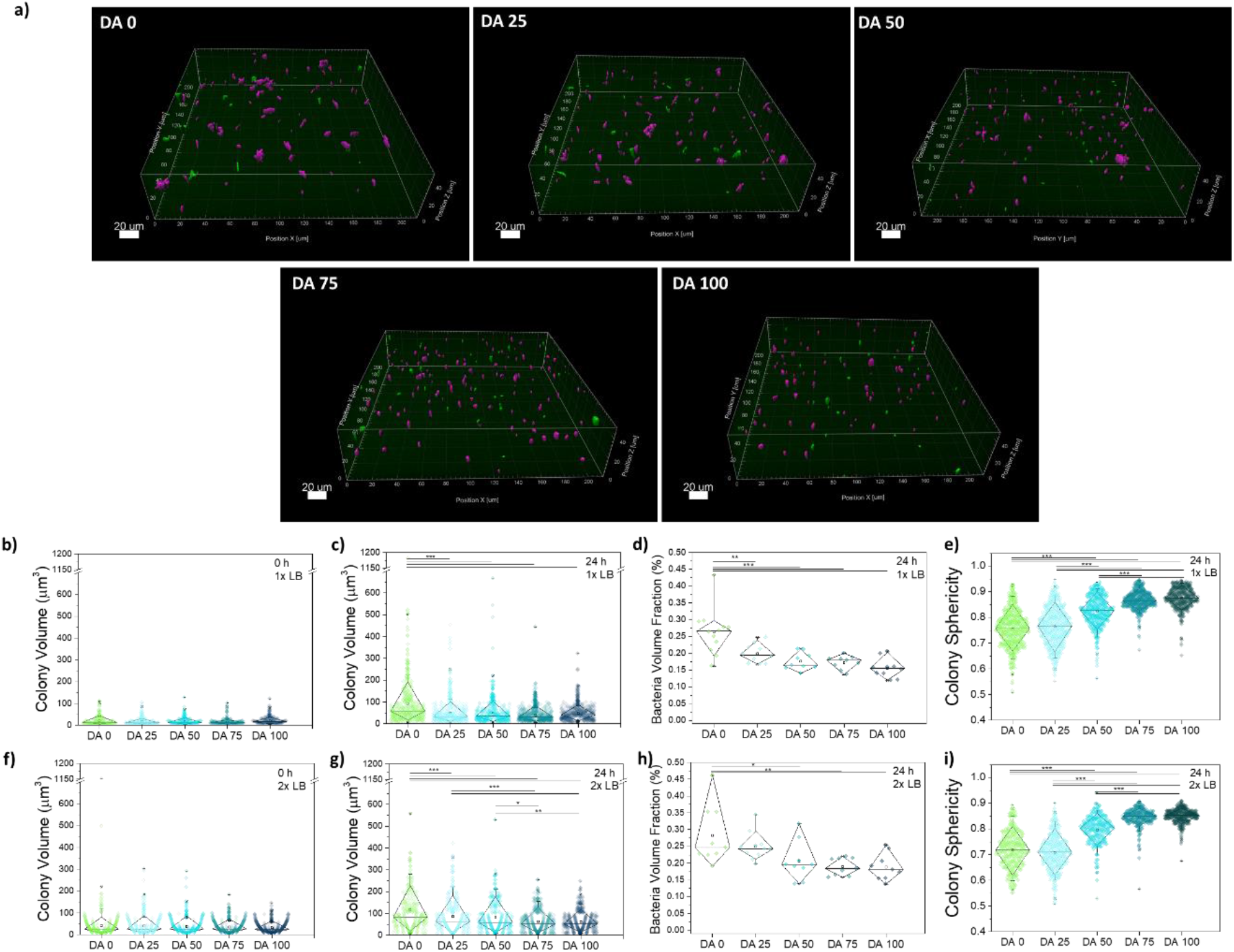
Bacteria colony growth inside hydrogels in microchannels (Figure S3b microchannel, filled with bacterial gel volume of 30 ul). a) Fluorescence image of 212.5 × 212.5 × 50.102 µm^3^ volumes of 1x LB concentration bacterial hydrogels producing iLOV after 24 h culture. The magenta-colored colonies do not touch the edges of the ROI, while the green colored colonies do. The magenta and green colored colonies were used for the quantification of the bacteria volume fraction (d,h). Only magenta-colored colonies were considered for the quantification of the volume (b,c,f,g) and sphericity (e,i) of individual colonies. For visualization surface masks obtained by IMARIS software were used. The scale bar corresponds to 20 μm; b) Colony volume distribution within different hydrogels of 1x LB concentration at 0 h and c) 24 h; d) Volume fraction of bacteria colonies within different hydrogels of 1x LB concentration at 24 h timepoint; e) The sphericity of the colonies within different hydrogels of 1x LB concentration at 24 h timepoint and f) Colony volume distribution within different hydrogels of 2x LB concentration at 0 h and g) 24 h; h) Volume fraction of bacteria colonies within different hydrogels of 2x LB concentration at 24 h timepoint and i) The sphericity of the colonies within different hydrogels of 2x LB concentration. (Diamond plots indicating 10 and 90 percentile values, whiskers indicating 5 and 95 percentile values, * p < 0.05, ** p < 0.01, *** p < 0.001)

**Figure S12.**
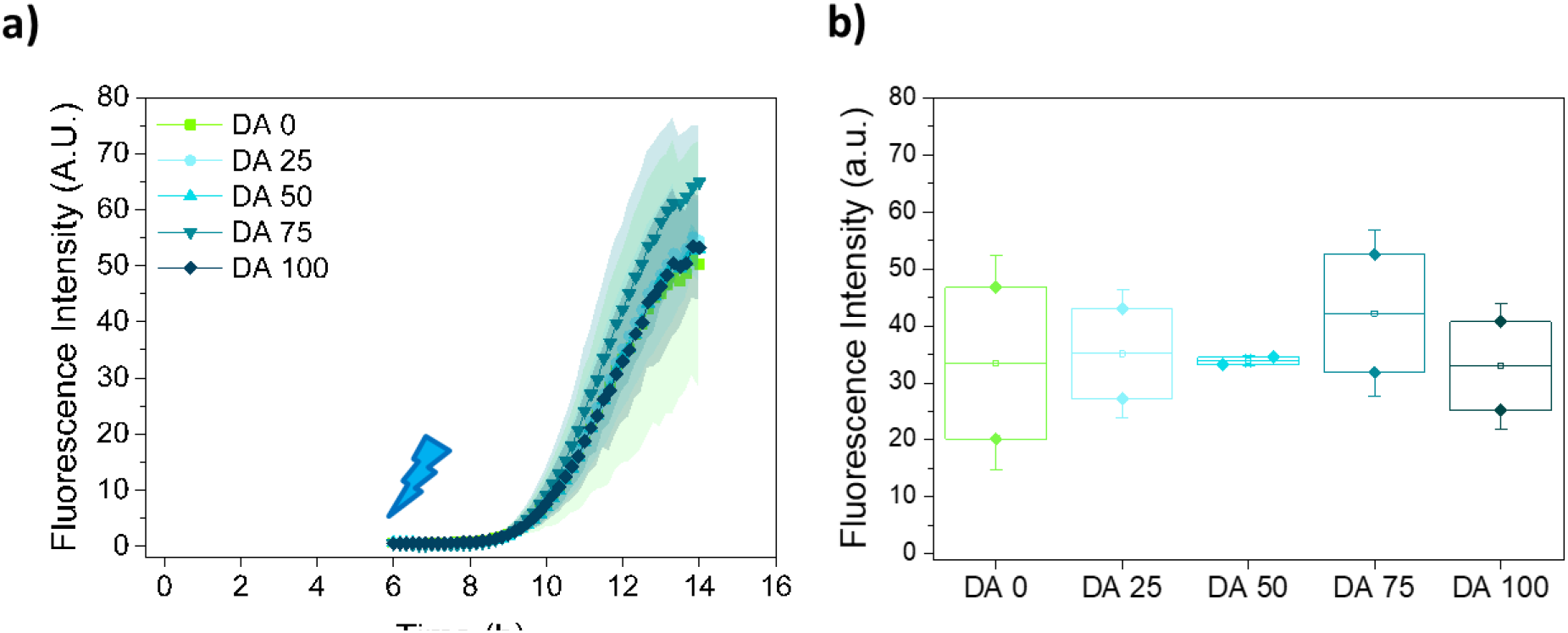
RFP production analysis. **a)** Normalized fluorescence intensity (mean ± standard deviation) associated to RFP production measured in bacteria loaded DA 0-100 hydrogels in microchannels (Figure S3b). The hydrogels were cultured for 6 h and then irradiated to induce protein production. **b)** Fluorescence intensity measured in the hydrogels at 12 h (6 h after the onset of irradiation). (N = 2, box represents 25 and 75 percentile values, whiskers indicate standard deviation)

### Different formats lead to different metabolic conditions but result in similar bacterial behavioral trends

When the gel-encapsulated bacteria were allowed to grow in micro-channels, confocal imaging at 0 h and 24 h timepoints indicated that colony volumes increased over time but were smaller (5 – 8x on average) compared to colony volumes in the micro-well format. The trend of colony volumes decreasing and sphericity increasing with increase in chemical cross-linking degree was still observed and was more pronounced when using medium with 2x concentration of nutrients, indicating that the hydrogels were imparting the same type of mechanical constraints in both microwell and micro-channel formats. Another notable aspect was that the fluorescence intensities within the cells were sufficient for quantification at 0 h and 24 h timepoints but not at earlier timepoints (3 h, 6 h), during which the highest rates of bacterial growth are expected to be occurring based on the data from Figure 3.

When RFP-expressing bacteria were encapsulated within the different types of gels and formed within the microchannels with one interface accessible to air, fluorescence appeared only to a depth of ∼ 1mm from that interface (**Figure S14**). This indicated that oxygen, which is required for RFP fluorophore formation, does not diffuse deeper into the gel, causing the bacteria to switch to anaerobic metabolic pathways for growth and sustenance, leading to smaller colony volumes and weaker protein production, as found by previous reports.^[3]^ Such oxygen diffusion limitations are not expected in the hydrogel films in the microwell since the silicone covering film is permeable to oxygen (Figure 3). Nevertheless, even under these conditions, trends of decreasing colony volumes, increasing sphericity and narrowing of distributions for both with increasing chemical cross-linking degrees were similar to the trends observed in the micro-well format. This indicated that the mechanical constraints imparted by the chemical cross-links influenced colony morphologies independent of metabolic variations.

**Figure S13.**
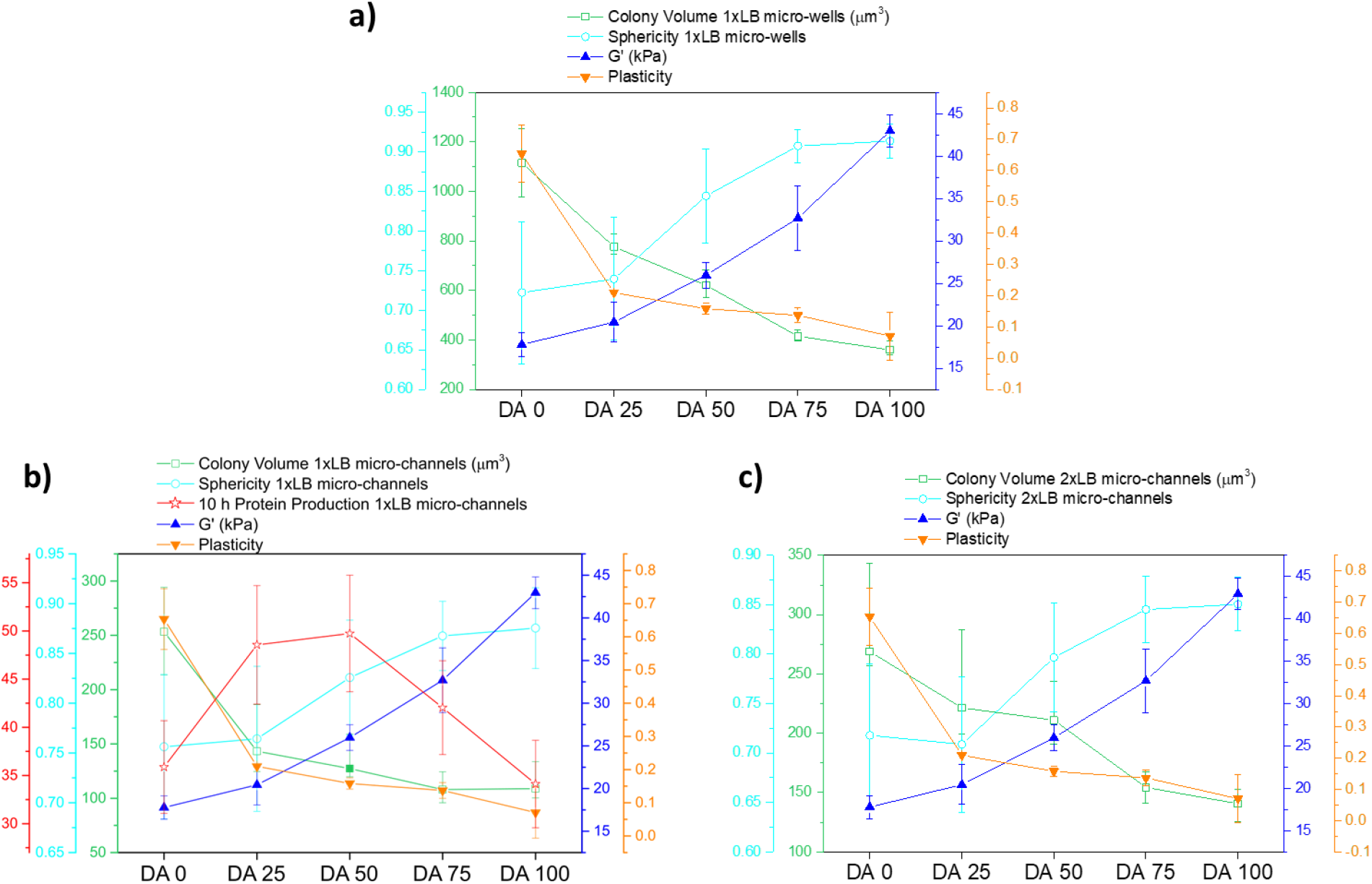
Comparative representation of hydrogel mechanical properties and the behavior of embedded bacteria colonies in DA 0-100 hydrogels. Mean value of shear storage modulus (from Figure 1b) and degree of plasticity (from Figure 1d), 95^th^ percentile of colony volume values and mean sphericity after 24 h, and mean value protein production after 10 h (from Figure 4d) within the indicated constructs were plotted. The error bars indicate the 93^rd^-97^th^ percentile value range for the colony volume data, for the rest, the error bars indicate standard deviation.

**Figure S14.**
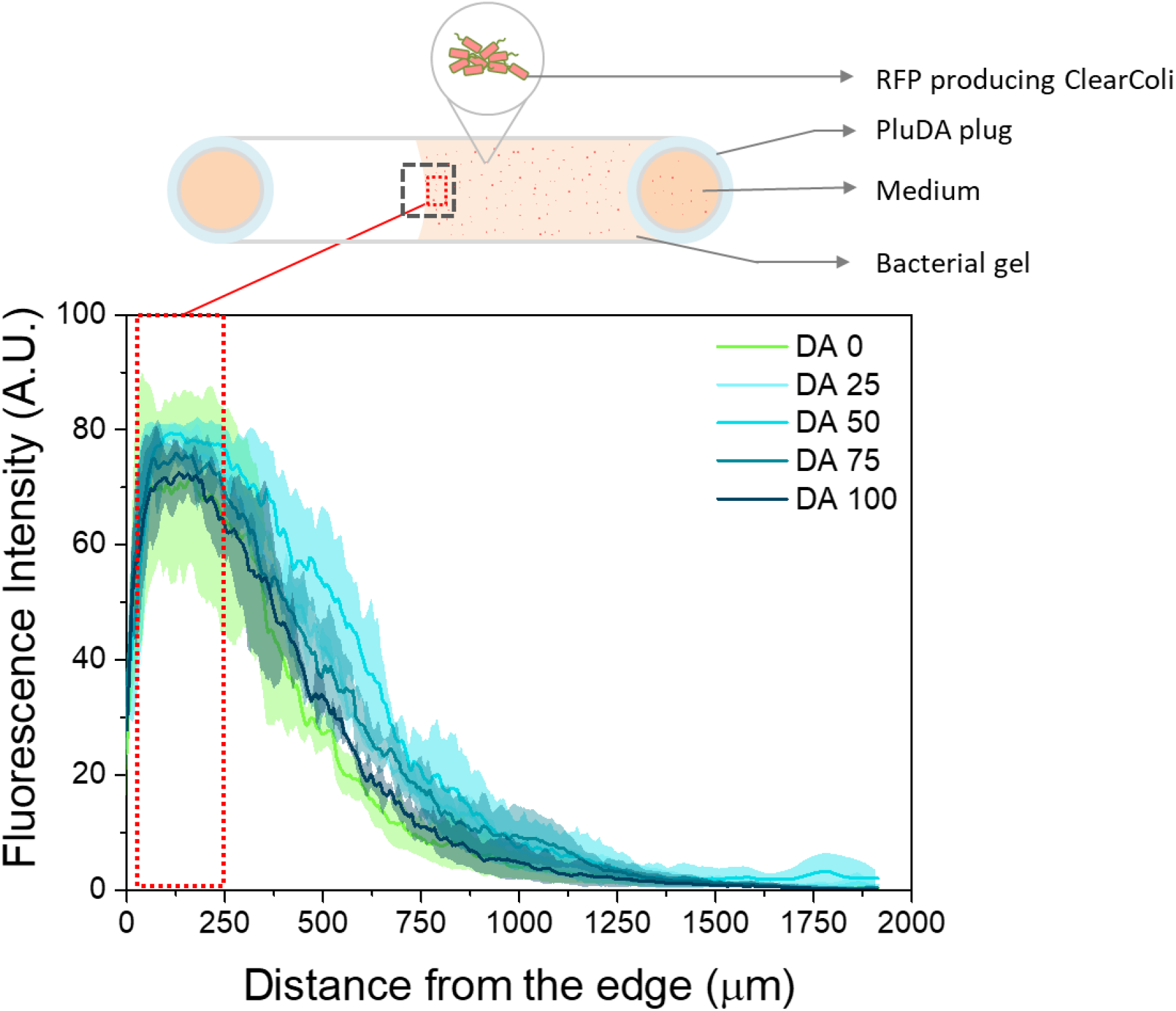
Fluorescence intensity profile in RFP producing bacteria hydrogels inside microchannels (Figure S3b) at t = 18 h. The fluorescence was measured from the edge of the hydrogel to the end of the channel along the channel longitudinal axes. The highlighted dotted box (200 × 600 µm^3^) indicates the region of the bacterial hydrogel considered for RFP fluorescence intensity measurement (Figure 4). (mean ± SD, N=3)

## Notes

### Competing Interest Statement

The authors have declared no competing interest.

